# Subcellular proteomics of dopamine neurons in the mouse brain reveals axonal enrichment of proteins encoded by Parkinson’s disease-linked genes

**DOI:** 10.1101/2021.06.01.446584

**Authors:** Benjamin D. Hobson, Se Joon Choi, Rajesh K. Soni, David Sulzer, Peter A. Sims

**Affiliations:** Department of Systems Biology, Columbia University Irving Medical Center, New York, NY 10032; Medical Scientist Training Program, Columbia University Irving Medical Center, New York, NY 10032; Department of Neurology, Columbia University Irving Medical Center, New York, NY 10032; Department of Psychiatry, Columbia University Irving Medical Center, New York, NY 10032; Department of Pharmacology, Columbia University Irving Medical Center, New York, NY 10032; Division of Molecular Therapeutics, New York State Psychiatric Institute, New York, NY 10032; Proteomics Shared Resource, Herbert Irving Comprehensive Cancer Center, Columbia University Irving Medical Center, New York, NY 10032; Department of Biochemistry & Molecular Biophysics, Columbia University Irving Medical Center, New York, NY 10032; Sulzberger Columbia Genome Center, Columbia University Irving Medical Center, New York, NY 10032; Aligning Science Across Parkinson’s (ASAP) Collaborative Research Network, Chevy Chase, MD

## Abstract

Dopaminergic neurons modulate neural circuits and behaviors via dopamine release from expansive, long range axonal projections. The elaborate cytoarchitecture of these neurons is embedded within complex brain tissue, making it difficult to access the neuronal proteome using conventional methods. Here, we demonstrate APEX2 proximity labeling within genetically targeted neurons in the mouse brain, enabling subcellular proteomics with cell type-specificity. By combining APEX2 biotinylation with mass spectrometry, we mapped the somatodendritic and axonal proteomes of midbrain dopaminergic neurons. Our dataset reveals the proteomic architecture underlying proteostasis, axonal metabolism, and neurotransmission in these neurons. We find a significant enrichment of proteins encoded by Parkinson’s disease-linked genes in striatal dopaminergic axons, including proteins with previously undescribed axonal localization. These proteomic datasets provide a resource for neuronal cell biology, and this approach can be readily adapted for study of other neural cell types.

## Introduction

Dopamine (DA) release from the axons of midbrain dopaminergic (DA) neurons provides important signals that regulate learning, motivation, and behavior. Given that dopaminergic dysfunction is linked to neuropsychiatric diseases including Parkinson’s disease (PD), schizophrenia, and drug addiction, there is considerable interest in deep molecular profiling of DA neurons in health and disease. Molecular profiling of striatal DA axons is of particular interest, since they may be the initial site of DA neuronal degeneration in PD (Burke & O’Malley, 2013), a point strongly supported by neuropathological comparison of dopaminergic axonal and cell body loss in PD patients (Kordower et al., 2013). Thus, the study of proteins in striatal DA axons and how they are altered in disease or disease models is a topic of intense investigation.

Despite their importance in behavior and disease, DA neurons are very few in number, with an estimated ~21,000 in the midbrain of C57BL/6 mice (Nelson et al., 1996). Thus, although proteomic profiling of midbrain tissue samples offers insight into PD pathophysiology (Jung et al., 2017; Petyuk et al., 2021), such studies do not analyze striatal DA axons, nor can they identify the cellular source of changes in protein levels. Most DA neuron-specific molecular profiling studies have focused on mRNA using translating ribosome affinity purification (TRAP) or single cell RNA-sequencing (scRNA-seq) (Agarwal et al., 2020; Brichta et al., 2015; Dougherty, 2017; D. J. Kramer et al., 2018; Poulin et al., 2014; Saunders et al., 2018; Tiklová et al., 2019). Although these methods have advanced our understanding of gene expression in DA neurons, there are important issues that are not addressed by the study of mRNA. First, mRNA levels do not always correlate with protein abundance, particularly for axonal proteins (Moritz et al., 2019). Second, transcriptomics cannot establish the localization or abundance of the encoded proteins within specific subcellular compartments, which is particularly important for DA neurons.

After exiting the midbrain, the axons of DA neurons travel within the medial forebrain bundle (MFB) to innervate forebrain structures. DA axonal arbors within the striatum are immense and highly complex: single DA neuron tracing has shown that DA axons can reach over 500,000 μm in total length (Matsuda et al., 2009). Although their axons constitute the major volume and energy demand for DA neurons (Bolam & Pissadaki, 2012; Matsuda et al., 2009; Pacelli et al., 2015), dopaminergic axons represent only a small fraction of tissue protein in the striatum. The ability to directly interrogate subcellular proteomes of DA neurons in native brain tissue would significantly enhance our understanding of DA neuronal biology.

To study the subcellular proteome of DA neuron in the mouse brain, we have adapted enzyme-catalyzed proximity labeling combined with mass spectrometry (MS)-based proteomics, a powerful approach for identifying protein interactions and/or localization in subcellular compartments (Hung et al., 2014; Loh et al., 2016; Rhee et al., 2013; Roux et al., 2012). For in-cell proximity labeling, a particularly efficient enzymatic approach uses the engineered ascorbate peroxidase APEX2 (Lam et al., 2015). APEX2 rapidly biotinylates proximal proteins in the presence of hydrogen peroxide (H_2_O_2_) and biotin-phenol (BP), generally requiring <1 minute of labeling in cell culture (Hung et al., 2016, 2017). While most proximity labeling studies have been conducted on cultured cells, APEX2 proximity labeling has also been demonstrated in live Drosophila tissues (C.-L. Chen et al., 2015), and recently in mouse heart (G. Liu et al., 2020). Selective expression of APEX2 enabled electron microscopy reconstructions of genetically targeted neurons in mice (Joesch et al., 2016; Q. Zhang et al., 2019), suggesting that APEX2 may be suitable for cell-type-specific proximity labeling and proteomics in live brain tissue.

Here, we employ APEX2-mediated biotin labeling in acute brain slices to study the subcellular proteome of DA neurons. Using a combination of cell type-specific APEX2 expression and proximity biotinylation in acutely prepared slices, we have characterized the somatodendritic and axonal proteomes of midbrain DA neurons. We show that the striatal axonal arbors contain nearly 90% of DA neuronal proteins accessible to cytoplasmic APEX2 labeling, providing robust coverage of proteins involved in axonal transport, dopamine transmission, and axonal metabolism. Of particular interest, we find that proteins encoded by DA neuron-enriched genes and PD-linked genes are preferentially localized in striatal axons, including novel axonal proteins. Together, our data establishes a proteomic architecture for DA neurons and identifies candidates for mechanistic follow-up studies of PD-relevant DA neuronal cell biology.

## Results

### APEX2 enables rapid, DA neuron-specific biotin labeling in acute brain slices

To express cytoplasmic APEX2 specifically in DA neurons, we employed an AAV5 viral vector containing the Cre-dependent construct developed by Joesch et al. (2016), which expresses a V5-tagged APEX2 fused to a nuclear export sequence (NES) (Wen et al., 1995). We injected AAV5-CAG-DIO-APEX2NES into the ventral midbrain (VM) of DAT^IRES-Cre^ mice (Bäckman et al., 2006) (**Figure 1a**). To confirm the cellular specificity and Cre-dependence of APEX2 expression, we injected the virus into DAT^IRES-Cre^ mice crossed with Ai9 tdTomato reporter mice (Madisen et al., 2010). Immunostaining against V5 (APEX2), RFP (tdTomato), and tyrosine hydroxylase (TH), a canonical DA neuron marker, demonstrated specific expression of APEX2 in DA neurons (**Figure 1b**). We found that ~97% of V5-APEX2^+^ neurons were also double-positive for tdTomato and TH, while V5-APEX2 expression was never detected in non-dopaminergic (TH^-^/tdTomato^-^) neurons (**Figure 1 – figure supplement 1a-b**). All three markers displayed intense staining throughout the DA neuronal cytoplasm, including dendrites in the VM and axonal projections in the striatum (**Figure 1b**). Thus, injection of Cre-dependent AAV-APEX2NES (hereafter referred to as APEX2) into the VM of DAT^IRES-Cre^ mice leads to robust expression of APEX2 throughout the DA neuronal cytoplasm.

**Figure 1:**
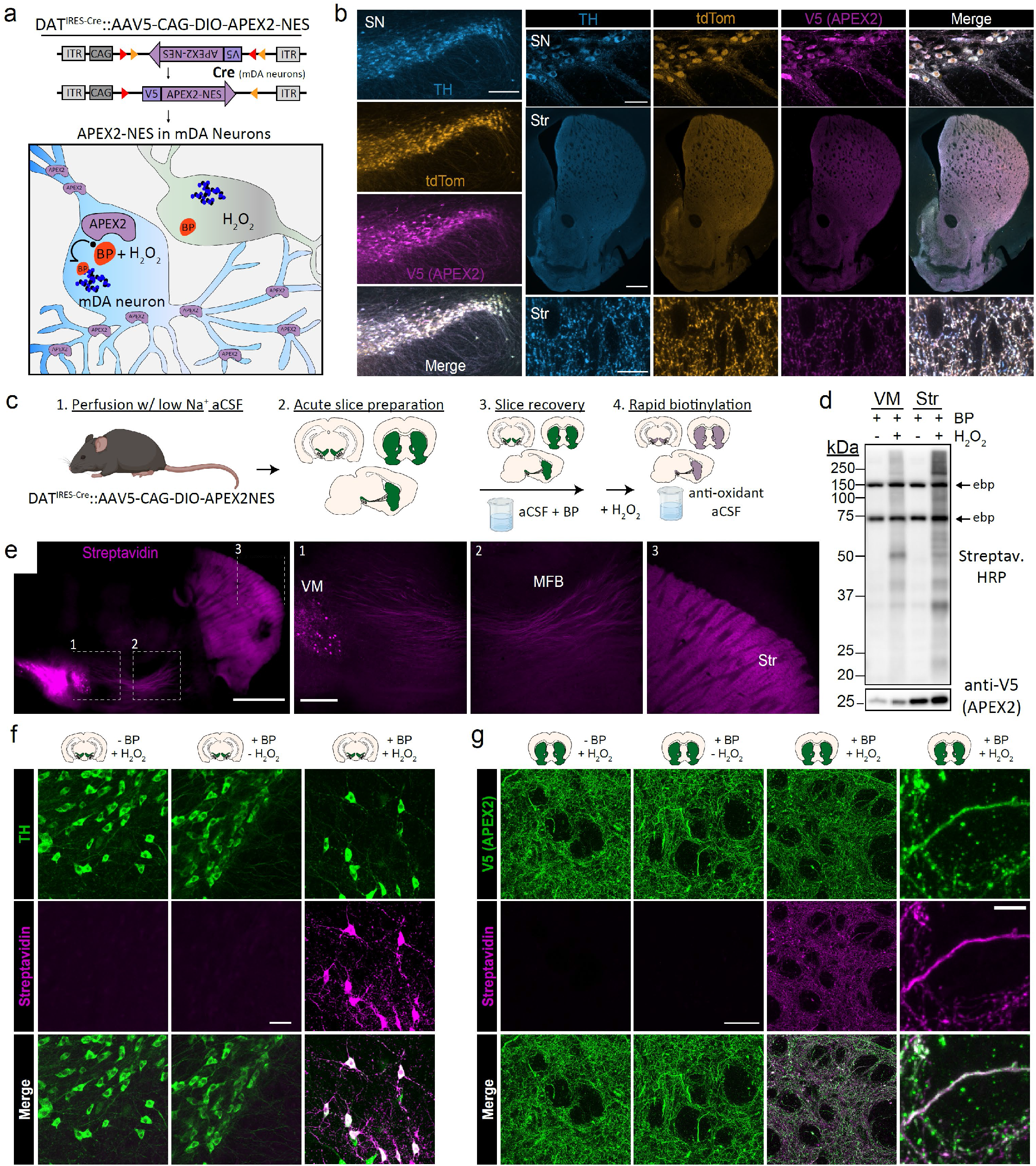
Cre-dependent viral expression of cytoplasmic APEX2 and mDA neuron-specific biotinylation in acute brain slices. (**a**) Schematic depicting viral expression strategy: Cre-dependent, cytoplasmic APEX2 expressing AAV (AAV5-CAG-DIO-APEX2-NES) is injected into the midbrain of DAT^IRES-Cre^ mice for mDA neuron-specific APEX2 labeling. (**b**) Immunostaining of mDA neurons in DAT^IRES-Cre^/Ai9^tdTomato^ mice injected with AAV5-CAG-DIO-APEX2-NES. Anti-TH, anti-RFP (tdTomato), and anti-V5 (APEX2) all display diffuse localization throughout somatic, dendritic, and axonal cytoplasm. *Left*, substantia nigra, scale bar: 200 μm, *upper right:* substantia nigra at high power, scale bar: 50 μm, *middle right:* dorsal and ventral striatum, scale bar: 500 μm, *lower right*: dorsal striatum at high power, scale bar: 10 μm. (**c**) Schematic depicting APEX2 labeling procedure in acute brain slices. Slice are incubated for one hour with 0.5 mM biotin phenol prior to labeling with 1 mM hydrogen peroxide for 3 minutes. (**d**) *Upper*: Western blotting of ventral midbrain and striatal slice lysates with streptavidin-horseradish peroxidase in the presence of biotin phenol with or without hydrogen peroxide (H_2_O_2_). Endogenously biotinylated proteins (ebp) are noted in all lanes at ~75 and ~150 kDa. *Lower*: Same as above but with anti-V5 (APEX2). (**e**) Streptavidin-AlexaFluor647 staining of sagittal slices after APEX2 labeling in mDA neurons. *Left*, sagittal slice at low power, scale bar: 1 mm. Insets indicated in white dashed lines, *right:* insets of the ventral midbrain, medial forebrain bundle, and striatum, scale bar: 250 μm. (**f**) Streptavidin-AlexaFluor647 staining of coronal midbrain slices after APEX2 labeling in mDA neurons, scale bar: 50 μm. Labeling requires both biotin phenol and hydrogen peroxide. (**g**) Same as (**f**) but with fields of striatal slices. First three columns, scale bar: 50 μm. Far right panels at high magnification, scale bar: 5 μm. *Abbreviations:* (aCSF) artificial cerebrospinal fluid, (BP) biotin phenol, (H_2_O_2_) hydrogen peroxide, (HRP) horseradish peroxidase, (MFB) medial forebrain bundle, (NES) nuclear export sequence, (TH) tyrosine hydroxylase, (SN) substantia nigra, (Str) striatum, (VM) ventral midbrain.

As shown in Drosophila (C.-L. Chen et al., 2015), APEX2 labeling can be performed in living tissue in the presence of H_2_O_2_ and BP. We chose to conduct labeling in acute brain slice preparations under conditions that preserve the integrity of neurons and severed axons, as shown by electrophysiological recordings and stable levels of evoked DA release for at least six hours (Hernandez et al., 2012). Typical sets of coronal or sagittal slices are shown in **Figure 1 – figure supplement 2a** and our workflow is summarized in **Figure 1c**. Key steps in the protocol include: 1) transcardial perfusion with low-sodium cutting solution, which acts to preserve neuronal integrity in slices from adult mice (Ting et al., 2014) and removes catalase-rich blood, 2) incubation of slices with BP in oxygenated artificial cerebrospinal fluid (aCSF) during the slice recovery period, 3) rapid labeling with H_2_O_2_ in aCSF, and 4) rapid quenching by transferring slices to antioxidant aCSF. Because the slices are far thicker (300 μm) than monolayer cell cultures, we fixed, cleared, and stained them with fluorescent anti-V5 (APEX2) and streptavidin after biotin labeling in 1 mM H_2_O_2_ for 1-5 minutes (**Figure 1 – figure supplement 2b**). We found that biotinylation was detectable at all time points, although streptavidin labeling appeared weaker within the center of slices, suggesting incomplete penetration of BP and/or H_2_O_2_. Rather than targeting a specific organelle or protein complex, our goal in this work was to broadly label the entire cytoplasm of DA neurons. Therefore, we chose 3 minutes of H_2_O_2_ exposure for downstream applications, which provided sufficient labeling for proteomics while limiting H_2_O_2_ exposure.

Western blotting of slices treated with BP and H_2_O_2_ showed broad biotinylation patterns in the midbrain and striatum (**Figure 1d**), consistent with labeling of somatodendritic and axonal proteins, respectively. Fluorescent streptavidin staining of sagittal slices after labeling and fixation revealed DA neuron-specific labeling throughout the VM, MFB, and striatum (**Figure 1e**). We confirmed that both BP and H_2_O_2_ are required for APEX2-mediated biotinylation in DA neuronal soma/dendrites and axons (**Figure 1f-g**). Confocal imaging of striatal slices revealed a dense, intricate staining pattern consistent with the cytoarchitecture of DA axons (**Figure 1g**). Biotin labeling co-localized with V5-APEX2^+^ DA axons but not with surrounding soma or myelin tracts, since APEX-generated BP radicals do not cross membranes (Rhee et al., 2013). These results show that APEX2 can rapidly and specifically label the somatodendritic and axonal compartments of DA neurons in acute brain slices.

### Proteomic profiling of subcellular compartments in DA neurons

We next used DA neuron-specific biotin labeling in slices to perform proteomic profiling of these neurons with subcellular resolution. After biotin labeling and quenching, we rapidly dissected and froze the VM, MFB, and striatum of sagittal slices for downstream biotinylated protein enrichment and liquid chromatography tandem mass spectrometry (LC-MS/MS) (**Figure 2a**). After lysis and protein precipitation to remove free biotin, we immunoprecipitated (IP) proteins from each region with streptavidin beads. To control for non-specific binding and potential labeling by endogenous tissue peroxidases, we prepared slices from DAT^IRES-Cre^ mice without APEX2 (APEX2^-^, no virus control) and treated them identically to APEX2^+^ slices at all stages of the protocol. Streptavidin horseradish peroxidase (HRP) blotting of captured proteins showed endogenously biotinylated carboxylase proteins at ~75 and ~150 kDa in all samples, while biotinylated proteins across a wide range of molecular weights were found only in APEX2^+^ samples (**Figure 2b**). Quantification of streptavidin HRP signal revealed that approximately 87% of DA neuronal APEX2 biotinylation is found within the striatum, 9% in the VM, and 4% in the MFB (**Figure 2d**). These results demonstrate that the majority of DA neuronal proteins are found within striatal axons.

**Figure 2:**
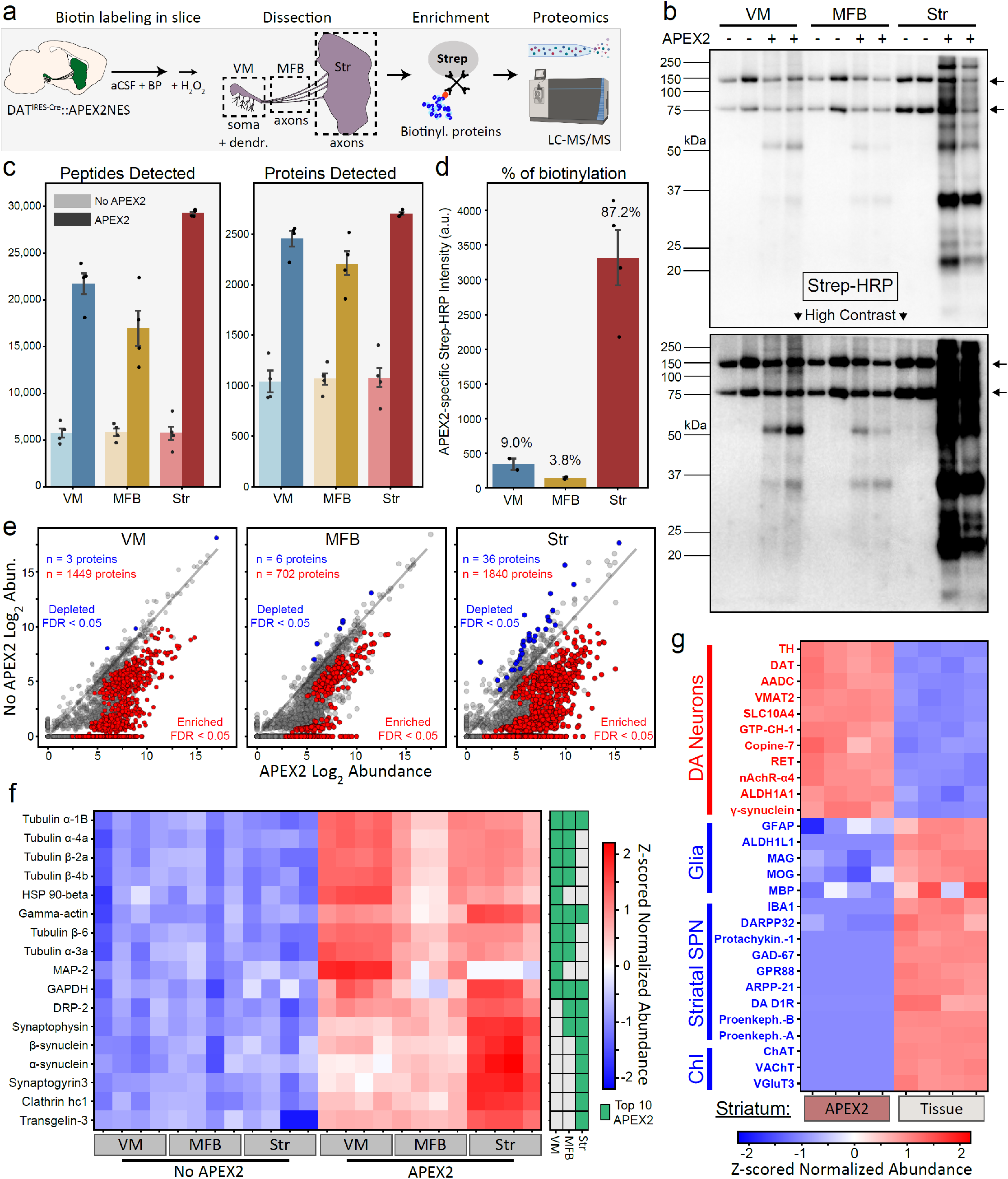
APEX2 proximity labeling proteomics in mDA neurons. (**a**) Schematic depicting APEX2 proximity labeling proteomics in mDA neurons. Slices are labeled, rapidly quenched, dissected, and flash frozen. Frozen tissues are lysed and precipitated to remove free biotin, after which re-solubilized tissue proteins are subjected to streptavidin bead immunoprecipitation to enrich biotinylated proteins. On-bead digestion produces peptides which are quantified by liquid chromatography with tandem mass spectrometry (LC-MS/MS). (**b**) Streptavidin-HRP western blotting of streptavidin IPs from tissue dissections of the indicated regions (duplicate lanes are biological replicates). Each lane contains proteins eluted from 5% of the streptavidin beads for each biological replicate (single mouse/region). Arrows on the right indicate prominent bands in APEX2^+^ and APEX2^-^ samples of all regions, which represent endogenous biotinylated carboxylase proteins at ~75 and ~150 kDa. The majority of APEX2-specific biotinylation is found in the striatum, but specific labeling is present in both VM and MFB at high contrast (*lower*). (**c**) Mean ± SEM of peptides and proteins detected per biological replicate of APEX2^-^ or APEX2^+^ streptavidin IPs of indicated regions (n = 4 each). See **Figure 2 – source data 2** for raw label-free quantification intensity values of peptides and proteins for all samples used in this study. (**d**) Quantification of streptavidin-HRP reactivity in APEX2^+^ streptavidin IPs, related to panel (**b**). After subtraction of endogenously biotinylated protein signal within each lane, the APEX2-specific streptavidin-HRP intensity for each APEX2^+^ sample was determined by subtracting the average APEX2^-^ lane intensity for the same region. Mean ± SEM normalized streptavidin-HRP intensity is plotted for each region (n=2 for VM, n=2 for MFB, n=4 for Striatum). The percentage of APEX2-specific biotinylation found in each region are denoted above the bars. (**e**) Log-log abundance plots of APEX2^-^ vs. APEX2^+^ streptavidin IP samples for the indicated regions. Axes represents the average log_2_(normalized intensity + 1) of n=4 biological replicates for each sample type. Proteins significantly enriched or depleted from APEX2^+^ streptavidin IP samples are colored in red or blue, respectively. False discovery rate (FDR) represents q values from Benjamini-Hochberg procedure on Welch’s (unequal variance) t-test. See **Figure 2 – source data 4** for complete results of APEX2^+^ vs. APEX2^-^ comparisons. (**f**) Heatmap of Z-scores for protein abundances for the union of the top 10 most abundant proteins enriched in APEX2^+^ vs. APEX2^-^ differential expression analysis from panel (e). Each column represents a biological replicate (n=4) of APEX2^-^ and APEX2^+^ streptavidin IP samples in the indicated regions. The green color bar on the right indicates whether a given protein was in the top 10 of each region. (**g**) Heatmap of Z-scores for protein abundances for markers of midbrain DA neurons, glia, striatal spiny projection neurons (SPN), and cholinergic interneurons (ChI). Each column represents a biological replicate (n=4) of bulk striatal tissue or APEX2^+^ streptavidin IP samples. See **Figure 2 – source data 5** for complete results of bulk tissue vs. APEX2^+^ comparisons. *Abbreviations:* (aCSF) artificial cerebrospinal fluid, (BP) biotin phenol, (H_2_O_2_) hydrogen peroxide, (HRP) horseradish peroxidase, (MFB) medial forebrain bundle, (Str) striatum, (VM) ventral midbrain. See **Figure 2 – source data 1** for list of protein abbreviations in (**f-g**).

After on-bead tryptic digestion, we conducted label-free quantitative proteomics for single-mouse biological replicates of VM, MFB, and striatum IP samples. Using data-independent acquisition (DIA), we quantified between 15,000-30,000 peptides representing 2,100-2,600 proteins per APEX2^+^ sample, while only ~5,000 peptides representing ~1,000 proteins were detected in APEX2^-^ samples (**Figure 2c**). Using the same LC-MS/MS workflow to analyze the bulk tissue proteome of VM and striatum slices, we found that approximately 45% of proteins quantified in these samples were also detected in APEX2^+^ samples (**Figure 2 – figure supplement 1a**). More than 84% of proteins in both bulk tissue and APEX2^+^ IP samples were identified based on quantification of multiple peptides (**Figure 2 – figure supplement 1b**), and protein abundances of biological replicate APEX2^+^ IP samples were highly correlated (**Figure 2 – figure supplement 1c**; Pearson’s r=0.93-0.94 for striatum, r=0.89-0.92 for VM, and r= 0.82-0.84 for MFB). Thus, even for specific subcellular compartments of small populations (~21,000 DA neurons), the high efficiency of APEX2 labeling enables highly reproducible cell type-specific proteomics from individual mice. We also conducted bulk tissue proteomics on VM and striatum slices that were immediately frozen or subjected to APEX2 labeling procedures (**Figure 2 – figure supplement 2a**). We found thousands of differentially expressed proteins when comparing VM vs. striatum slices, but no statistically significant differences when comparing acute vs. rested slices (**Figure 2 – figure supplement 2b-c**). We also found that incubation of slices with 0.5 mM BP for one hour had no effect on spontaneous action potential frequency in DA neurons (**Figure 2 – figure supplement 2d-e**). These data demonstrate that acute slice preparation and labeling procedures do not compromise DA neuronal function or significantly distort brain tissue proteomes.

To identify APEX2-dependent proteins captured by streptavidin IP, we directly compared APEX2^+^ to APEX2^-^ control samples (**Figure 2e**). The number of proteins enriched in APEX2^+^ samples scaled with the fraction of biotinylation derived from each region (see **Figure 2b-d**), with 1449 proteins for VM, 702 proteins for MFB, and 1840 proteins for striatal samples (FDR < 0.05, Welch’s unequal variance t-test with Benjamini-Hochberg correction). Although proteins such as endogenously biotinylated carboxylases (e.g., pyruvate carboxylase, propionyl-CoA carboxylase) and other non-specific binders (e.g., myelin basic protein, proteolipid protein 1) were abundant in all samples, they were not enriched in APEX2^+^ samples. The most abundant APEX2-specific proteins within each region displayed considerable overlap across regions (**Figure 2f**), with 17 common proteins derived from the top 10 proteins in each region (VM, MFB, and striatum). Although cytoskeletal proteins such as actin and tubulin subunits were highly abundant in APEX2^+^ samples from all three regions, the most abundant APEX2-specific proteins within each region also included proteins enriched in specific subcellular compartments. For example, the somatodendritic protein MAP-2 (microtubule-associated protein 2) was in the top 10 only for VM samples (**Figure 2f**), while proteins involved in synaptic vesicle fusion and endocytosis were in the top 10 only for MFB or striatum samples (e.g., synaptophysin, synaptogyrin-3, alpha-synuclein). Compared to bulk striatal tissue, APEX2^+^ striatal samples show significant enrichment of dopaminergic proteins such as TH, dopamine transporter (DAT), aromatic-/-acid (dopa) decarboxylase (AADC), and vesicular monoamine transporter 2 (VMAT2) (**Figure 2g**). Meanwhile, proteins specific to astrocytes, microglia, oligodendrocytes, striatal spiny projection neurons (SPNs), and cholinergic interneurons (ChI) were either not detected or were significantly depleted from striatal APEX2^+^ samples (**Figure 2g**). Thus, mass spectrometry-based quantification of APEX2-enriched proteins enables proteomic profiling of the dopaminergic neuronal compartments contained within each region.

### Somatodendritic vs. axonal enrichment of proteins involved in diverse cellular functions

To further establish the compartment-specificity of APEX2-specific proteins across regions, we first examined the abundance of the microtubule binding proteins MAP-2 and tau, which are known to be enriched in somatodendritic and axonal compartments, respectively. Consistent with previous work (Okabe & Hirokawa, 1989; Papasozomenos et al., 1985), we found that MAP-2 was highly abundant in the VM and steadily decreased in the MFB and striatum, while tau was most abundant in striatal samples (**Figure 3a**). It should be noted that although we refer to VM samples as somatodendritic, these samples will necessarily include axons exiting the midbrain as well as somatically synthesized axonal proteins. Accordingly, the difference in abundance between striatal and VM samples was dramatically greater for MAP-2 compared to tau (**Figure 3a**). Thus, for our downstream analysis of somatodendritic and axonal compartments, we used the MFB samples only for filtering and focused primarily on VM and striatum samples.

**Figure 3:**
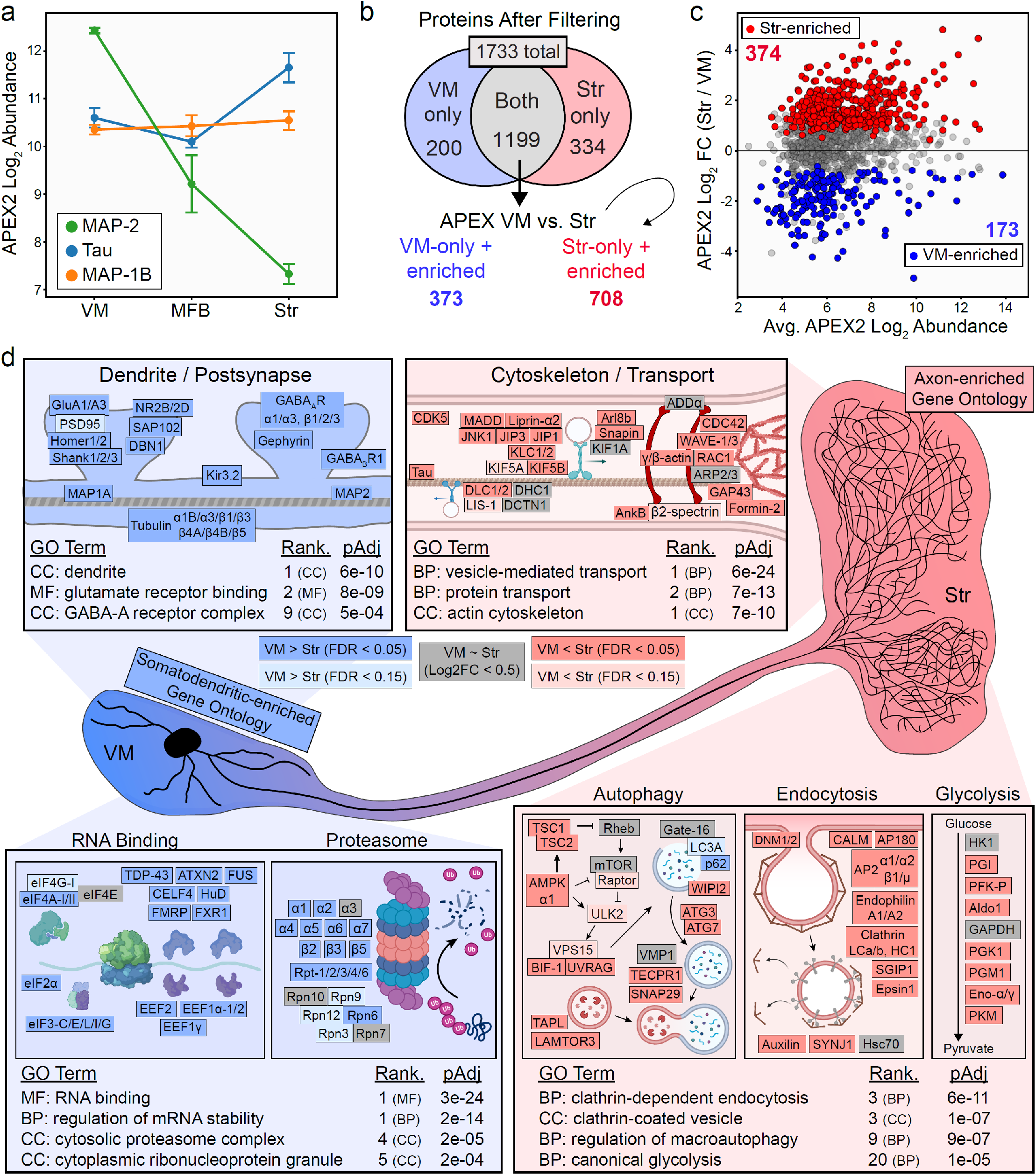
Axonal and somatodendritic proteomics reveals gene ontologies enriched in subcellular compartments. **(a)** APEX2 proteomics data for microtubule-associated proteins MAP-2, Tau, and MAP-1B. Mean ± SEM of the protein abundances, as log2(normalized intensity + 1), are shown for n=4 biological replicates of APEX2^+^ streptavidin IP samples in the indicated regions. **(b)** Schematic depicting proteins remaining after filtering. For complete filtering workflow, see **Methods** and **Figure 3 – figure supplement 1**. Proteins present in both VM and Str after filtering were further compared by differential expression—see panel (**c**). Proteins present only in VM or Str, plus those enriched in VM vs. Str differential expression analysis, were used for subsequent GO Analysis. **(c)** Differential expression comparison of VM vs. Str APEX2^+^ streptavidin IP samples. Proteins colored red or blue had a false discovery rate (FDR) < 0.05 after Benjamini-Hochberg corrected p-values from Welch’s (unequal variance) t-test. See **Figure 3 – source data 2** for complete results and summary of proteins before and after filtering. **(d)** Gene ontology analysis (*Enrichr*) of VM- and Str-enriched proteins (373 and 708, respectively, see panel b). Selected GO Terms are listed along with adjusted p-values and adjusted p-value rank for each GO Term category (Cellular Component, Molecular Function, Biological Process). Canonical and representative proteins from the ontologies are shown. Every protein depicted is present in the filtered proteomics data of VM, Str, or both. Colors indicate significant (dark red/blue) enrichment, near-significant enrichment (light red/blue), or similar levels between VM and Str (gray). Slashes indicate separate proteins (e.g., DLC1/2 represents both DLC1 and DLC2). See **Figure 3 – source data 3** for complete GO summary. *Abbreviations:* (MAP-2) Microtubule-associated protein 2, (Tau) Microtubule-associated protein tau, (MAP-1B) Microtubule-associated protein 1B, (MFB) medial forebrain bundle, (Str) striatum, (VM) ventral midbrain. See **Figure 3 – source data 1** for list of protein abbreviations in (**d**).

To filter the APEX2^+^ VM and striatum proteomics data, we took advantage of publicly available scRNA-seq data from the mouse ventral midbrain and striatum (Saunders et al., 2018). After identification of high-confidence DA neuron profiles, we used the mRNA expression distribution from scRNA-seq to establish a conservative lower bound for considering a gene as expressed in DA neurons (**Figure 3 – figure supplement 1a**, see **Methods**). The vast majority of proteins in the APEX2 data were encoded by genes expressed in DA neurons, with >93.0% of all detected proteins and >97.4% of APEX2-enriched proteins encoded by genes above our scRNA-seq threshold (**Figure 3 – figure supplement 1b-c**). We removed proteins that were not significantly enriched in APEX2+ > APEX2-samples, proteins encoded by genes below our DA neuron scRNA-seq threshold, and proteins that did not show evidence of APEX2-specificity in at least two regions (**Figure 3 – figure supplement 1d**, see **Methods** for complete description). For comparison between VM and striatal samples, we retained the union of proteins passing filters in VM and striatal samples. Out of 1,733 total proteins, 200 proteins passed filtering only in VM samples, 334 only in striatal samples, and 1,199 both (**Figure 3b**). Hierarchical clustering of these 1,199 overlapping proteins clearly segregated VM APEX2^+^ and Str APEX2^+^ samples from each other and from all APEX2^-^ samples (**Figure 3 – figure supplement 2**), and direct comparison between APEX2^+^ samples revealed 173 and 374 proteins with greater relative abundance in VM or striatal samples, respectively (**Figure 3c**). Manual examination of VM- and Str-enriched protein clusters revealed striatal enrichment of synaptic vesicle proteins (e.g., Synaptotagmin-1 (Syt-1), Synaptophysin, SNAP25, VMAT2) and VM enrichment of postsynaptic scaffolding proteins (e.g., Homer-2, Shanks1-3) (**Figure 3 – figure supplement 2**).

We first sought to confirm that cytoplasmic APEX2 labeling does not enrich proteins within membrane-enclosed structures. Amongst the 1733 total proteins passing filter in either VM or striatum, we found significant enrichment of gene ontology (GO) terms related only to nuclear and mitochondrial outer membranes (**Figure 3 – figure supplement 3a-b**), but not nuclear inner membrane, nuclear lamina, nuclear matrix, mitochondrial inner membrane, or mitochondrial matrix. These results confirm the membrane impermeability of BP radicals and the integrity of organellar membranes in the acute brain slices.

To assess the relative enrichment of functionally related proteins within the somatodendritic and axonal compartments of DA neurons, we conducted GO analysis of proteins that either passed filtering in only one region or were significantly enriched in that region in the APEX2^+^ vs. APEX2^-^ comparison (**Figure 3b-c**, 373 proteins for VM, 708 for striatum). First, we analyzed subcellular localization GO terms curated by the COMPARTMENTS resource (Binder et al., 2014). We found that the top GO terms over-represented amongst VM-enriched proteins included ‘Postsynapse’, ‘Somatodendritic compartment’, ‘Dendrite’, and ‘Postsynaptic density’ (**Figure 3 – figure supplement 3c**), while those over-represented amongst striatum-enriched proteins included ‘presynapse’, ‘axon’, ‘synaptic vesicle’, and ‘axon terminus’ (**Figure 3 – figure supplement 3d**). Given the clear segregation of axonal/pre-synaptic and dendritic/post-synaptic terms from the COMPARTMENTS resource in striatum- and VM-enriched proteins, respectively, we analyzed the VM- and Str-enriched proteins using the GO Consortium resource (Gene Ontology Consortium, 2021). Similar to the COMPARTMENTS analysis, we found that GO terms related to postsynaptic function were over-represented in VM-enriched proteins, with terms such as ‘dendrite’, ‘glutamate receptor binding’, and ‘GABA-A receptor complex’. Canonical proteins associated with these terms included subunits and scaffolding proteins for AMPA, NMDA, GABA-A, and GABA-B receptors (**Figure 3d**). GO terms such as ‘RNA binding’ and ‘regulation of mRNA stability’ were also over-represented in VM-enriched proteins (**Figure 3d**), consistent with the somatodendritic compartment being the major site of protein synthesis in neurons. Despite a clear role for the ubiquitin proteasome system in axons (Korhonen & Lindholm, 2004), we found over-representation of GO terms such as ‘cytosolic proteasome complex’ in VM-enriched proteins, with nearly all 20S subunits and most 19S subunits showing higher relative abundance in VM APEX2^+^ samples (**Figure 3d**).

The enrichment of protein synthesis and degradation machinery within the somatodendritic compartment of DA neurons underscores the importance of cytoskeletal transport systems in axonal protein homeostasis (Maday et al., 2014; Roy, 2014). Accordingly, GO terms such as ‘vesicle-mediated transport’ and ‘protein transport’ were over-represented in striatum-enriched proteins. Striatum-enriched proteins in these ontologies included kinesin and dynein subunits, cargo adaptor proteins, and upstream kinases involved in transport regulation (**Figure 3d**). In addition to microtubule-based transport proteins, proteins related to the axonal actin cytoskeleton were also over-represented in striatum APEX2^+^ samples. These included actin subunits themselves, GTPases and other regulators of actin nucleation, and proteins that link actin rings in the distal axon (Leterrier et al., 2017; Xu et al., 2013) (**Figure 3d**). We also found that proteins involved in clathrin-dependent endocytosis were uniformly enriched in striatum APEX2^+^ samples, consistent with high rates of synaptic vesicle recycling in DA neurons related to their tonic activity. Given the intense energetic demands placed on DA axons (Pissadaki & Bolam, 2013), we were intrigued to find an over-representation of glycolytic enzymes in Str-enriched proteins: 7 out of 9 glycolytic enzymes show higher relative abundance in striatum compared to VM (**Figure 3d**). These results suggest that glycolysis may be especially important in dopaminergic axonal metabolism, consistent with glycolysis supporting axonal transport and presynaptic function in other neurons (Ashrafi et al., 2017; Hinckelmann et al., 2016; Jang et al., 2016; Zala et al., 2013). We note that striatum APEX2^+^ samples also showed extensive enrichment of GO terms related to presynaptic function and synaptic vesicle proteins, as detailed below (see **Figure 3 - source data 3** for a complete GO analysis summary).

While anterograde transport is critical for delivery of new proteins to distal axons, it is likely that both local autophagy and retrograde transport contribute to protein clearance in DA axons. Many autophagosomes formed in distal axons are transported back to the soma prior to fusion with lysosomes (Maday et al., 2012), although other findings implicate local autophagic degradation of damaged mitochondria in axons (Ashrafi et al., 2014). Indeed, we previously showed that macroautophagy regulates presynaptic structure and function in DA axons (Hernandez et al., 2012). Consistent with these results, we found that GO terms related to autophagy were over-represented in Str-enriched proteins (**Figure 3d**). Autophagy-related proteins enriched in striatum APEX2^+^ samples included kinases involved in upstream regulation of autophagy, membrane proteins involved in autophagosome maturation, and membrane proteins involved in lysosomal membrane fusion. Among the autophagy-related proteins present in striatal axons, vacuole membrane protein 1 (VMP1) was recently shown to be critical for survival and axonal integrity in DA neurons (Wang et al., 2021). Our results suggest that VMP1 may regulate autophagy in the axons as well as soma of DA neurons, highlighting the ability of APEX2 proteomics to elucidate the distribution of proteins with unknown localization. Collectively, these results establish a foundation of protein localization that underlies diverse cellular functions within the somatodendritic and axonal compartments of DA neurons.

### The dopaminergic presynaptic proteome via APEX2 labeling in striatal slices and synaptosomes

DA neuron-specific APEX2 labeling in striatal slices provides a convenient means to isolate proteins from dopaminergic axons, but acute slice preparation may not be practical for all laboratories. To further study the presynaptic proteome of dopaminergic axons and provide an alternative to the slice labeling procedure, we conducted APEX2 labeling in synaptosomes prepared from the striatum of DAT^IRES-Cre^ mice expressing APEX2NES (**Figure 4a**). Synaptosomes are resealed nerve terminals formed by liquid shearing forces during homogenization of brain tissue in isotonic sucrose buffer (Gray & Whittaker, 1962; Whittaker, 1993). After a brief, low-speed centrifugation to remove heavy cellular debris and nuclei, the majority of synaptosomes can be rapidly recovered in the pellet from moderate-speed centrifugation known as the P2 fraction. Although the P2 fraction also contains myelin and free mitochondria, we reasoned that these contaminants would not interfere with APEX2 labeling within resealed dopaminergic nerve terminals. After isolation and washing of the P2 fraction, reagents were added directly to synaptosomes under typical *in vitro* APEX2 labeling conditions (30 minutes incubation with 0.5 mM BP followed by 60 seconds of 1 mM H_2_O_2_). Similar to the slice procedure, streptavidin-HRP western blotting of striatal P2 lysates showed APEX2-dependent protein biotinylation across a wide range of molecular weights (**Figure 4b**). Staining of labeled P2 samples with fluorescent streptavidin revealed deposition of biotin within TH^+^/V5-APEX2^+^ particles approximately ~1 μm in diameter, consistent with APEX2 labeling within dopaminergic synaptosomes (**Figure 4b**). We analyzed APEX2-enriched proteins from striatal synaptosomes by mass spectrometry. As before, we controlled for non-specific binding by conducting all labeling and protein purification procedures on striatal synaptosomes from mice with and without APEX2 expression in DA neurons.

**Figure 4:**
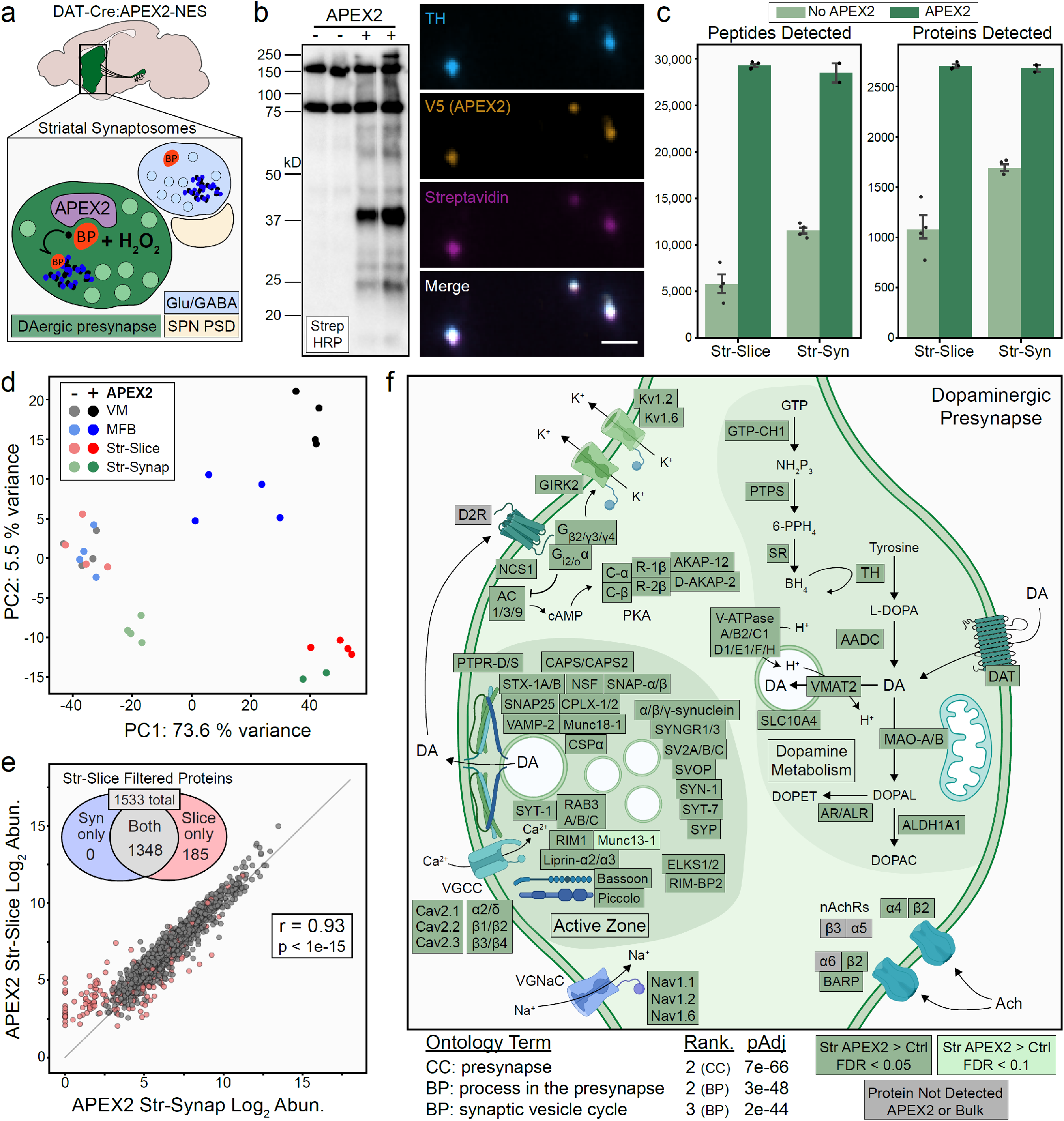
APEX2 labeling in synaptosomes and dopaminergic presynaptic proteome. (**a**) Schematic depicting APEX2 labeling in dopaminergic striatal synaptosomes. A crude synaptosome fraction (P2) is rapidly prepared from the striatum, washed, incubated with biotin phenol (0.5 mM for 30 minutes), labeled with 1 mM H_2_O_2_ for 60 seconds, quenched with antioxidants and sodium azide, centrifuged, and flash frozen as a pellet for downstream streptavidin IP and proteomics (see **Methods**). (**b**) *Left:* Streptavidin HRP western blot of proteins captured by streptavidin IP from striatal synaptosomes of DAT-Cre mice with or without expression of APEX2-NES. *Right:* Immunostaining of synaptosomes from DAT-Cre:APEX2^+^ striatum. Synaptosomes were bound to poly-lysine coated coverslips during biotin phenol incubation, washed, and fixed after H_2_O_2_ treatment. Scale bar: 2 μm. (**c**) Mean ± SEM of peptides and proteins detected per biological replicate of APEX2^-^ or APEX2^+^ streptavidin IPs of indicated regions. Str-slice data are the same as in same as in Figure 2c (n=4 each for APEX2^-^ /APEX2^+^). n=4 for APEX2^-^ Str-Syn, n=2 for APEX2^+^ Str-Syn. (**d**) Principal components analysis of all APEX2^-^ and APEX2^+^ biological replicates for the 1733 proteins present either in VM or Str after filtering (see Figure 3b). PC1 captures 73.6% of the variance and is dominated by APEX2^+^ vs. APEX2^-^ samples, while PC2 captures 5.% of the variance and stratifies regional APEX2^+^ samples. APEX2^+^ Str-Syn samples are highly similar to APEX2^+^ Str-Slice samples. (**e**) Log-log abundance plots of APEX2^+^ streptavidin IP samples for the indicated regions. Axes represents the average log_2_(normalized intensity + 1) for each sample type (n=2 and n=4 biological replicates for Str-Synaptosomes and Str-Slice, respectively). Out of 1533 Str-Slice APEX2^+^ filtered proteins (see **Figure 3 – figure supplement 1d**), 1348 show concordant enrichment in Str-Syn APEX2^+^ vs. APEX2^-^ comparisons (log_2_ FC > 0 and FDR < 0.15). These 1348 proteins were retained for Gene Ontology analysis (panel f). Accordingly, Str-Syn and Str-Slice APEX2^+^ samples show robust Pearson correlation (r = 0.93, p value < 1e-15). (**f**) Gene ontology analysis (*SynGO*) of Str-Slice- and Str-Syn-enriched proteins (1348 proteins, see panel (**e**). Selected GO Terms are listed along with adjusted p-values and adjusted p-value rank for each GO Term category (Cellular Component, Biological Process). Canonical and representative proteins from the ontologies are shown. Proteins depicted are either present in the filtered Str-Slice proteomics data (FDR < 0.05, APEX2 vs. Control, dark green), nearly missed the Str-Slice significance thresholds (FDR < 0.1, APEX2 vs. Control, light green), or were not detected in any APEX2 or bulk striatal tissue samples (grey). See **Figure 4 – source data 2** for complete GO summary. *Abbreviations:* (SPN PSD) Spiny projection neuron post-synaptic density, (TH) Tyrosine hydroxylase, (MFB) medial forebrain bundle, (Str) striatum, (VM) ventral midbrain. See **Figure 4 – source data 1** for complete list of protein and metabolite abbreviations in (**f**).

Coverage of both peptides and proteins in APEX2^+^ synaptosome samples was remarkably similar to the APEX2^+^ striatal slice samples (**Figure 4c**), although non-specific binding was slightly higher in APEX2^-^ synaptosome samples. Principal component analysis revealed tight co-segregation of APEX2^+^ striatal slice and synaptosome samples apart from all other samples, suggesting these APEX2^+^ proteomes are highly similar (**Figure 4d**). Indeed, the correlation between APEX2+ striatal slice and synaptosome samples was comparable to that of biological replicates within each sample type (Pearson’s r = 0.93, p < 1e-15) (**Figure 4e**). Out of 1,533 proteins that passed filtering in striatal slices, 1,348 of these showed enrichment in APEX2^+^ vs. APEX2^-^ synaptosomes (**Figure 4e**).

Proteins involved in DA metabolism were significantly enriched in both slice and synaptosome APEX2^+^ samples, including canonical DA synthesis, release, and reuptake proteins (TH, AADC, VMAT2, and DAT) as well as enzymes involved in tetrahydrobiopterin synthesis (GTP cyclohydrolase I, 6-pyruvoyltetrahydropterin synthase, and sepiapterin reductase) and DA degradation (monoamine oxidase A/B, retinaldehyde dehydrogenase 1, aldose reductase, and aldehyde reductase) (**Figure 4f**). Thus, all of the protein machinery required for DA neurotransmission and metabolism is present within dopaminergic axonal boutons, including monoamine oxidases that metabolize DA and contribute to oxidative phosphorylation via presynaptic mitochondria (Graves et al., 2020). We also found significant enrichment of proteins that contribute to DA vesicular uptake, including most subunits of the vesicular ATPase and SLC10A4, an orphan transporter found on monoaminergic synaptic vesicles that contributes to axonal DA homeostasis (Larhammar et al., 2015) (**Figure 4f**).

Striatal DA release is powerfully modulated by nicotinic receptors and DA D2 receptors on dopaminergic axons (Sulzer et al., 2016). Consistent with pharmacological studies (Exley & Cragg, 2008), we found that β2 and α4 nicotinic receptor subunits were enriched by APEX2 in striatal slices and synaptosomes (**Figure 4f**). Despite a wealth of functional evidence supporting their presence on DA axons, the DA D2 receptor as well as β3, α5, and α6 nicotinic receptor subunits were notably absent from our dataset. Since the vast majority of APEX2 labeling occurs on tyrosine residues (Kim et al., 2018; Udeshi et al., 2017), some membrane proteins may not be labeled if they lack cytoplasm-facing tyrosine residues accessible to BP radicals. However, the absence of these nicotinic receptor subunits and the DA D2 receptor is likely related to mass spectrometry, since none of these proteins were detected in any bulk striatal tissue sample (**Figure 4f**, grey boxes). Thus, the absence of a protein in our dataset does not necessarily indicate absence in DA axons, especially when other functionally related proteins are enriched. For example, we found significant APEX2 enrichment of β-anchoring and -regulatory protein (BARP), a protein recently shown to interact with and modulate α6/β2/β3 nicotinic receptors on dopaminergic axons (Gu et al., 2019) (**Figure 4f**). Similarly, we find significant APEX2 enrichment of D2 receptor-interacting proteins (e.g., G protein subunits and neuronal calcium sensor NCS-1) and downstream effectors (e.g., adenylyl cyclase, PKA subunits, and GIRK2) known to function within DA neurons (Dragicevic et al., 2014). We also found significant enrichment of voltage gated potassium channels Kv1.2 and Kv1.6, consistent with previous work on Kv1 channels as downstream effectors of presynaptic D2 receptor function (Martel et al., 2011) (**Figure 4f**).

We next focused on the dopaminergic presynapse, using the SynGO resource (Koopmans et al., 2019) to analyze the 1,348 proteins enriched in APEX2^+^ striatal slice and synaptosome samples. Ontology terms such as ‘presynapse’, ‘synaptic vesicle’, and ‘presynaptic active zone’ were all highly over-represented amongst APEX2-enriched proteins (**Figure 4f** and **Figure 4 – figure supplement 1a**). Consistent with recent work on the molecular architecture of striatal DA release sites, APEX2 labeling of striatal and synaptosome samples significantly enriched active zone proteins RIM1, Bassoon, Liprin-α2 and -α3, Munc13-1, ELKS1/2, and RIM-BP2 (**Figure 4f**), although ELKS1/2 and RIM-BP2 are not essential for DA release (Banerjee, Imig, et al., 2020; C. Liu et al., 2018). Coverage of synaptic vesicle trafficking and fusion proteins was extensive, with significant enrichment of all major integral synaptic vesicle proteins (SV2A-C, CSPα, Synaptogyrin1-3, Synaptophysin, Synapsin-1, RAB3A-C, VAMP-2), target SNAREs (Syntaxin 1A/B, SNAP25), and functionally related proteins (NSF, SNAP-α/β, Munc18-1, Complexin1/2, and α/β/γ-synuclein). We also observed a significant enrichment of CAPS1 and CAPS2, proteins that prime synaptic vesicles in hippocampal neurons (Jockusch et al., 2007) and regulate catecholamine loading into large dense-core vesicles in adrenal chromaffin cells (Speidel et al., 2005) but whose function is to our knowledge unexplored in DA neurons. Notably, we find significant enrichment of Synaptotagmins-1 and −7 (Syt-1/Syt-7) (**Figure 4f**). These data support the recent finding that Syt-1 is the major fast Ca^2+^ sensor for synchronous DA release (Banerjee, Lee, et al., 2020). In addition to the established role of Syt-7 in somatodendritic DA release and the observation of its presence in DA axon terminals (Delignat-Lavaud et al., 2021; Mendez et al., 2011), our findings support Syt-7 as a candidate protein for asynchronous axonal DA release (Banerjee, Lee, et al., 2020). These data highlight candidate proteins for further investigation and provide a proteomic architecture of the dopaminergic presynapse that is highly consistent with recent functional studies (Banerjee, Imig, et al., 2020; Banerjee, Lee, et al., 2020; C. Liu et al., 2018).

Somewhat unexpectedly, we also found significant APEX2 enrichment of proteins with SynGO annotations for ontology terms related to postsynaptic function (**Figure 4 – figure supplement 1a**). Many of these proteins were annotated for both pre-and post-synaptic function, including proteins such as DAT and VMAT2 (**Figure 4 – figure supplement 1b**). Although these proteins are present within dopaminergic dendrites (Nirenberg et al., 1996a, 1996b), they are clearly derived from dopaminergic axons in our striatal APEX2^+^ samples. Similarly, many proteins with postsynaptic annotations were functionally related to actin dynamics, PKA or MTOR signaling, or other functions not exclusively related to postsynaptic function. Other proteins bearing only postsynaptic SynGO annotations, such as the Netrin receptor DCC and metabotropic glutamate receptor mGluR1, have functional evidence supporting their localization on DA axons (Reynolds et al., 2018; H. Zhang & Sulzer, 2003). Nonetheless, 63 proteins with clear roles in dendritic spines or mRNA translation were enriched by APEX2 in both slice and synaptosome samples (**Figure 4 – figure supplement 1b**). These proteins were significantly lower in abundance than proteins with presynaptic annotations (**Figure 4 – figure supplement 1c**). Although we cannot rule out the possibility of low levels of post-synaptic contamination, the significant depletion of proteins highly expressed by glia and striatal neurons (**Figure 2e**) demonstrates the absence of extensive cross-membrane labeling. Furthermore, we observed no enrichment of DARPP-32 and Spinophilin (**Figure 4 – figure supplement 2a**), proteins highly abundant within striatal dendritic spines (Allen et al., 1997; Blom et al., 2013; Greengard et al., 1999; Smith et al., 1999).

APEX2 enrichment in all regions can provide additional evidence of axonal localization for proteins often found in the post-synapse. For example, GABA receptors are typically present on dendrites, but recent electrochemical and electrophysiological studies have demonstrated the presence of both GABA-A and GABA-B receptors on DA axons (P. F. Kramer et al., 2020; Lopes et al., 2019; Schmitz et al., 2002). We found that GABA-A receptor subunits and scaffolding protein gephyrin were most abundant in VM APEX2^+^ samples, but still higher in APEX2^+^ than APEX2^-^ samples from both MFB and striatum (**Figure 4 – figure supplement 2b**). Similar results were obtained for GABA-B receptor subunits, the PDZ scaffold Mupp1 (Balasubramanian et al., 2007), and the effector protein GIRK2 (**Figure 4 – figure supplement 2c**). Although many comparisons did not pass the FDR correction in MFB comparisons, likely due to substantially lower proteomic depth in these samples, APEX2 enrichment in both MFB and striatal samples supports axonal transport and localization of these proteins.

Axonal APEX2 proteomics allowed us to identify proteins and protein complexes that are previously undescribed in striatal DA axons. The cytosolic chaperonin T-complex protein 1-ring complex (TRiC), or chaperonin containing T-complex (CCT) is an oligomeric complex that promotes folding of newly synthesized polypeptides, suppresses aggregation of huntingtin in Huntington’s disease, and can inhibit assembly of α-synuclein amyloid fibrils (Lopez et al., 2015; Sot et al., 2017; Tam et al., 2009). Recent work has implicated specific TRiC/CCT subunits in the regulation of axonal transport in cortical neurons (X.-Q. Chen et al., 2018; Zhao et al., 2016), but axonal localization of TRiC subunits is largely undescribed. We detected seven out of eight TRiC subunits in our APEX2 proteomics data, six of which showed strong evidence of axonal localization (**Figure 4 – figure supplement 2d**). These results suggest that the recently described interaction between CCT5/CCTε, CDK5, and tau (X.-Q. Chen et al., 2018) may regulate retrograde transport in DA neurons. Given that TRiC/CCT can regulate α-synuclein aggregation (Sot et al., 2017), future research on TRiC/CCT function in dopaminergic axons is warranted.

### Subcellular localization of proteins encoded by DA neuron-enriched and PD-linked genes

Genetic analysis can identify mutations that cause familial PD or variants linked to sporadic PD risk, but not whether the relevant genes are expressed in DA neurons. Although TRAP and scRNA-seq provide significant insight into DA neuronal gene expression, these techniques do not address the subcellular localization of the encoded proteins. We leveraged our APEX2 proteomics data to interrogate proteins encoded by genes with high DA neuron-specificity or by genes linked to PD via Mendelian inheritance or genome-wide association studies (GWAS) (**Figure 5a**). We re-analyzed published scRNA-seq data (Saunders et al., 2018) (**Figure 5 – figure supplement 1**), and identified 64 genes with >8-fold higher expression in DA neurons compared to all other midbrain cells (see **Methods**). 55 proteins encoded by these genes were present in the filtered APEX2 proteomics data (**Figure 5 – source data 1**). For human genes linked to familial PD, Atypical Parkinsonism, and Dystonia-Parkinsonism (Marras et al., 2016), we identified 15 proteins encoded by orthologous mouse genes in the filtered APEX2 proteomics data (out of 26 genes, **Figure 5 – source data 2**). For human genes nearest to PD-linked single nucleotide polymorphisms (SNPs) in the most recent GWAS meta-analysis (Nalls et al., 2019), we identified 86 orthologous mouse genes and 17 proteins encoded by these genes in the filtered APEX2 proteomics data (**Figure 5 – source data 2**).

**Figure 5:**
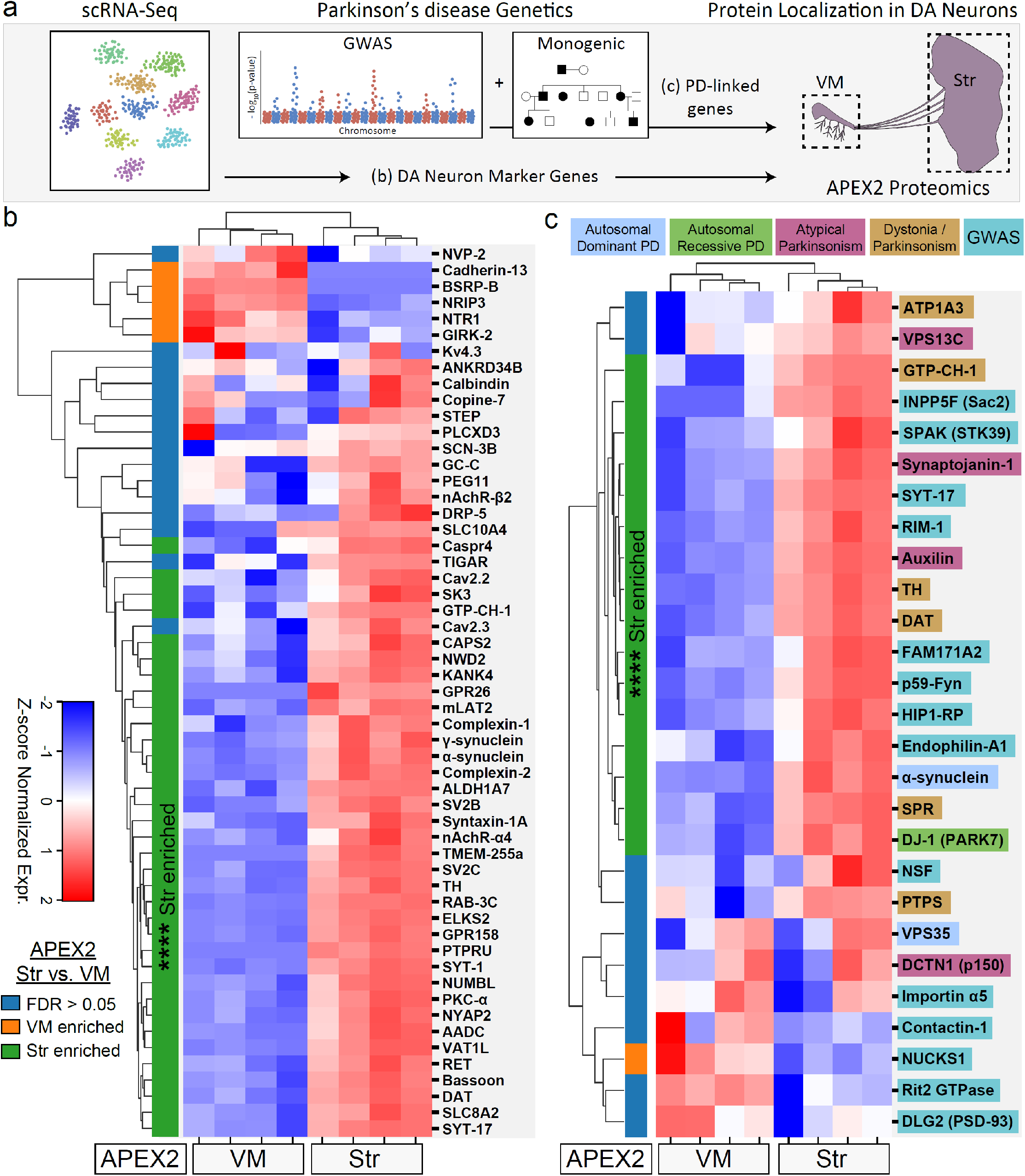
Subcellular proteomic analysis of proteins encoded by DA neuron marker genes and PD-linked genes. (**a**) Schematic depicting analysis workflow for integrating APEX2 proteomics data with scRNA-seq and genetic data. Proteins encoded by top DA neuron marker genes from mouse scRNA-seq data (Saunders et al., 2018) and human PD-linked gene orthologs (Marras et al., 2016; Nalls et al., 2019) are analyzed for subcellular protein localization in the APEX2 proteomics data. (**b**) Clustered heatmap of Z-scores for abundances of proteins encoded by the top 55 DA neuron marker genes present in the filtered APEX2 proteomics data. Each column represents a biological replicate (n=4) of APEX2^+^ streptavidin IP samples from the VM or striatum. The color bar on the left indicates whether a given protein was enriched in the VM (orange) or striatum (green) in differential expression analysis between APEX2^+^ IP samples (FDR < 0.05 after Benjamini-Hochberg corrected p-values from Welch’s t-test). **** indicates p = 4.8e-13, hypergeometric test for Str-enriched proteins (x=35) amongst the top 55 DA neuron markers present in the filtered proteomics data (1733 proteins). See **Figure 5 – source data 1** for summary of DA neuron marker genes and protein abbreviations. (**c**) Clustered heatmap of Z-scores for abundances of proteins present in the filtered APEX2 proteomics data and encoded by genes linked to hereditary PD/Parkinsonism or to PD via GWAS studies as indicated. Each column represents a biological replicate (n=4) of APEX2^+^ streptavidin IP samples from the VM or striatum. The color bar on the left indicates whether a given protein was enriched in the VM (orange) or striatum (green) in differential expression analysis between APEX2^+^ IP samples (FDR < 0.05 after Benjamini-Hochberg corrected p-values from Welch’s t-test). **** indicates p = 2.8e-06, hypergeometric test for Str-enriched proteins (x=17) amongst the 29 PD/Parkinsonism-linked genes present in the filtered proteomics data (1733 proteins). See **Figure 5 – source data 2** for list of human genes, orthologous mouse genes, and protein abbreviations.

Only a minority of proteins encoded by DA neuron-enriched and PD-linked genes were preferentially localized to the somatodendritic compartment of DA neurons (**Figure 5b-c**). In contrast, a significant majority of proteins encoded by DA neuron-enriched genes (35 out of 55, p = 5e-13, hypergeometric test, **Figure 5b**) and PD-linked genes (17 out of 29, p = 3e-06, hypergeometric test, **Figure 5c**) were preferentially localized to dopaminergic axons (FDR < 0.05 for APEX2^+^ striatal vs. VM samples). DA neuron-enriched genes encoding striatal APEX2-enriched proteins included many canonical synaptic vesicle and active zone proteins mentioned above (e.g., Bassoon, Syt-1, Complexin-1/2, SV2B/C, RAB3C, CAPS2), highlighting the strikingly high proportion of DA neuronal gene expression dedicated to presynaptic function (**Figure 5b**). Accordingly, many striatum-enriched proteins involved in vesicular release or endocytosis are encoded by genes linked to hereditary Atypical Parkinsonism (Synaptojanin-1, Auxilin) or to PD via GWAS (RIM-1, NSF, Endophilin-A1, and Sac2/INPP5F). Cao et al. (2020) recently demonstrated a role of Sac2/INPP5F in synaptic vesicle recycling, and our data confirm the presynaptic localization of endogenous Sac2/INPP5F in DA neurons (**Figure 5c**). These proteomic data provide strong support for genetic data implicating endocytic membrane trafficking pathways in PD risk (Bandres-Ciga et al., 2019). Another major group of APEX2^+^ striatum-enriched proteins were those involved in DA synthesis and transmission, many of which were DA neuron-specific (AADC), linked to hereditary Dystonia-Parkinsonism (6-pyruvoyltetrahydropterin synthase and sepiapterin reductase), or both (TH, DAT, and GTP cyclohydrolase I) (**Figure 5b-c**). These data suggest that dysfunction of tetrahydrobiopterin and/or DA synthesis pathways within dopaminergic axons may be central to the pathophysiology of these hereditary forms of Dystonia-Parkinsonism. Finally, we observed significant striatal enrichment of α-synuclein, VPS13C, and DJ-1 (**Figure 5c**), proteins encoded by familial PD genes. Collectively, these data are consistent with axonal dysfunction as a major contributor to DA neuronal degeneration in genetic PD (Burke & O’Malley, 2013).

In addition to the well-studied PD-related proteins noted above, many striatum-enriched proteins encoded by PD GWAS-linked genes have been largely undescribed in axonal function. We particularly highlight two candidates, SPAK/STK39 and Synaptotagmin-17 (Syt-17), both of which are encoded by the probable causal gene within their respective PD GWAS loci (Nalls et al., 2019). SPAK (sterile-20 (Ste20)-related proline-alanine-rich kinase) is a serine/threonine kinase that phosphorylates and modulates the activity of cation-chloride transporters, such as the Na+− K+–2Cl− cotransporter (NKCC1) and the K+–Cl− cotransporter KCC2 (Delpire & Austin, 2010; Geng et al., 2009). We find APEX2 enrichment of NKCC1, KCC2, SPAK, and upstream regulatory kinases in in VM, MFB, and striatal samples (**Figure 5 – figure supplement 2**). Given the role of NKCC1 and KCC2 in setting the chloride reversal potential (Kaila et al., 2014), recent demonstration of GABA-A receptor currents in DA axons (P. F. Kramer et al., 2020), and locomotor phenotype of SPAK knockout mice (Geng et al., 2010), these data warrant further study of SPAK function in dopaminergic axons. The axonal enrichment of Syt-17 is also of particular interest, given the linkage of human SYT17 to a PD risk locus and the enrichment of *Syt17* mRNA in DA neurons (**Figure 5b-c**). Syt-17 is an atypical synaptotagmin that does not bind calcium or participate in synaptic vesicle fusion (Ruhl et al., 2019) and has no established role in axons. Although hippocampal neurons from Syt-17 knockout mice display axonal growth defects, tagged Syt-17 is found in the Golgi complex of these cells and this phenotype appears to be mediated by deficits in vesicular trafficking (Ruhl et al., 2019). We are currently investigating the unknown function of SYT-17 in dopaminergic axons. Thus, our APEX2 proteomic dataset elucidates the subcellular localization of proteins encoded by PD-linked genes and highlights novel areas for further study of PD-relevant DA neuronal cell biology.

## Discussion

Our study demonstrates APEX2 labeling and mass spectrometry-based proteomics of axonal and somatodendritic compartments of DA neurons in the mouse brain. Thus, APEX2 labeling in acute brain slices provides a general approach for cell type-specific proteomics and/or proximity labeling proteomics in the mouse brain (see also, Dumrongprechachan et al., 2021), alongside promiscuous biotin ligase (BioID or TurboID)-catalyzed proximity labeling (Takano et al., 2020; Uezu et al., 2016) and incorporation of non-canonical amino acids via mutant tRNA synthetases (Alvarez-Castelao et al., 2017; Krogager et al., 2018). Each of these methods has advantages and disadvantages.

The major advantages of APEX2 are speed and efficiency: we were able to capture thousands of proteins from multiple subcellular compartments of a relatively rare neuronal population in individual mice (**Figure 2**). In comparison, BioID and TurboID labeling in the mouse brain typically requires over seven days and pooling tissue from many mice, precluding the use of individual mice as biological replicates (Takano et al., 2020; Uezu et al., 2016). The mutant tRNA synthetase methods label all synthesized proteins during 7 - 21 days of non-canonical amino acid administration, which complicates dynamic studies of specific protein complexes (Alvarez-Castelao et al., 2017; Krogager et al., 2018). In addition to its high efficiency, the speed of APEX2 labeling enables dynamic studies of protein complexes on a time scale of minutes (Lobingier et al., 2017), including during physiological responses in the mouse heart (G. Liu et al., 2020). Future studies might combine APEX2 labeling with optical and electrophysiological slice manipulations to interrogate rapid changes in neuronal physiology at the proteomic level. Beyond neurons, selective expression of APEX2-fusion proteins will broadly facilitate proteomic profiling of organelles and protein-protein interactions within a variety of genetically targeted brain cells.

APEX2 labeling in the mouse brain has several limitations compared to other aforementioned methods. First, the introduction of labeling reagents required preparation of acute brain slices or synaptosomes, which would preclude labeling during long-term behavioral or environmental manipulations (e.g., Alvarez-Castelao et al., 2017). Second, the high efficiency of APEX2 may lead to low levels of off-target protein labeling. Although we observed biotin labeling within morphologically intact DA axons (**Figure 1**) and significant depletion of proteins specific to striatal SPNs and glia (**Figure 2e**), we cannot completely exclude post-synaptic contamination. We propose at least four possible explanations for the detection of proteins with ‘postsynapse’ annotations in striatal APEX2 samples (**Figure 4 – figure supplement 1d**). First, it is possible that many canonical ‘post-synaptic’ proteins are also localized within DA axons. For example, we and others have shown that NMDA receptors are present on DA axons (Fortin et al., 2012; Schmitz et al., 2009), and most members of the membrane-associated guanylate kinase (PSD-93, PSD-95, SAP-102, SAP-97) and Shank families have been observed in axons and/or nerve terminals (Aoki et al., 2001; Arnold & Clapham, 1999; Halbedl et al., 2016; Muller et al., 1995). Similarly, the RNA binding- and translation-related proteins we observe may be derived from DA axons, given recent reports of translational machinery in mature axons (Hafner et al., 2019; Shigeoka et al., 2016; Younts et al., 2016). Second, it is possible that a small amount of APEX2 is transferred between DA axons and post-synaptic compartments during slice preparation and synaptosome homogenization. Third, it is possible that BP radicals directly cross damaged axonal and synaptosomal membranes during APEX2 labeling. Finally, it is possible that biotinylated proteins present only in APEX2^+^ samples provide surfaces for non-specific binding of non-biotinylated proteins. These mechanisms are not mutually exclusive, but appear to produce similar results in both slice and synaptosome labeling environments.

Although the slice procedure may not be accessible to all laboratories, APEX2 labeling in synaptosomes provides a simple and rapid alternative to access the presynaptic proteome of genetically targeted projection neurons. Somewhat surprisingly, we found that the APEX2 proteome of dopaminergic synaptosomes is largely comparable to that of the entire axonal arbor within the striatum (**Figure 4e**). We suspect this is due to the high density of boutons *en passant* along dopaminergic axons, which would provide a strong representation of the entire axonal proteome upon resealing as synaptosomes. Our data thus highlight the utility of the synaptosome sorting technique developed by Herzog and colleagues as a complementary approach for study of the presynaptic proteome (Biesemann et al., 2014; Pfeffer et al., 2020). Future work will determine whether the strong correlation between axonal and synaptosomal proteomes is a unique feature of DA neurons.

Mass spectrometry-based quantification of proteins provides significant advantages over immunohistochemical methods, especially for fine structures like dopaminergic axons. Suitable antibodies that provide high quality immunofluorescence in brain tissue are not available for a majority of the proteome, and off-target binding affects the reproducibility and interpretation of research findings (Weller, 2016). Many studies rely on overexpression of tagged proteins to establish localization, which can result in mis-localization and altered function. Our study demonstrates proximity labeling of endogenous proteins within subcellular compartments of genetically targeted neurons, and can thus be used for unbiased discovery as well as protein-targeted biochemical experiments.

While they represent the vast majority of neuronal volume and are critically important to DA biology, dopaminergic axons have remained largely inaccessible to proteomic study. Due to the relative immaturity, limited axonal complexity, and incomplete synaptic and hormonal inputs of cultured DA neurons, we chose to examine the DA neuronal proteome in native brain tissue of adult mice. Cytoplasmic labeling in dopaminergic axons captured a diverse range of cytosolic and membrane proteins involved in metabolism, protein transport, endolysosomal trafficking, synaptic transmission, and autophagy (**Figures 3–4**). Thus, our axonal proteomic dataset should be broadly useful to axonal and neuronal cell biologists.

The axonal enrichment of autophagy-related proteins is of particular importance to DA neurons, given that autophagy dysfunction is heavily implicated in PD pathophysiology (Wong & Cuervo, 2010) and associated with methamphetamine toxicity (Larsen et al., 2002). Due to the limitations of bulk striatal tissue-based protein measurements, our previous studies of autophagy in DA axons were limited to electrochemical measurement of DA release and morphological changes via electron microscopy (Hernandez et al., 2012). APEX2 labeling will enable future studies of protein biochemistry in DA axons, including proteins that are ubiquitously expressed and difficult to resolve using immunohistochemistry.

A majority of DA neuron-enriched genes encode proteins localized to axons (**Figure 5b**), suggesting that this compartment is central to the identity of DA neurons. These findings may be particularly important for understanding pathogenic mechanisms in PD. Importantly, we show that mutations linked to hereditary PD or parkinsonism are far more likely to be found in genes that encode axonal proteins (**Figure 5c**). Thus, the massive axons of DA neurons can be considered a double-edged sword: while they are required for DA release to support healthy brain function, they are susceptible to myriad environmental and genetic insults.

We note that the biggest risk factor for idiopathic PD is aging (Sulzer, 2007). It is tempting to speculate that the proteomic framework and cytoarchitecture of DA neurons in the mammalian brain evolved under positive selective pressure related to motor control, reward, and motivation, with little selective pressure related to the organism’s lifespan. Comparing the axonal proteome of DA neurons to that of other neurons spared from degeneration in PD (Surmeier et al., 2017) may identify distinguishing features that contribute to increased risk of axonal degeneration in PD. Thus, our study lays a proteomic foundation upon which future studies of neuronal cell biology and PD pathophysiology may build.

## Materials and Methods

### Animals

Adult male and female mice (6-12 months old) were used in all experiments. DAT^IRES-Cre^ mice (JAX #006660, RRID: IMSR_JAX:006660) and Ai9 mice (JAX #007909, RRID: IMSR_JAX:007909) were obtained from Jackson Laboratories (Bar Harbor, ME). Mice were housed on a 12-hour light/dark cycle with food and water available *ad/ibitum*. All experiments were conducted according to NIH guidelines and approved by the Institutional Animal Care and Use Committees of Columbia University and the New York State Psychiatric Institute.

### Plasmid and Virus

AAV-CAG-DIO-APEX2NES was a gift from Joshua Sanes (Addgene plasmid #79907;http://n2t.net/addgene:79907; RRID:Addgene 79907). AAV-CAG-DIO-APEX2NES was packaged into an AAV5 vector by Vector BioLabs (Malvern, PA). The final titer of the AAV5-CAG-DIO-APEX2NES preparation was 1 x 10^12^ GC/mL in PBS + 5% glycerol. To avoid freeze-thaw, single-use 10 μL aliquots were stored at −80°C.

### Viral Injection

All surgical procedures were approved by the Institutional Animal Care and Use Committee and the Department of Comparative Medicine at New York State Psychiatric Institute. Mice were anesthetized with 4% isoflurane. Animals were transferred onto a Kopf Stereotaxic apparatus and maintained under isoflurane anesthesia (1–2%). After hair removal and sterilization of the scalp using chlorhexidine and ethanol, a midline incision was made. Bregma and Lambda coordinates were determined, and minor adjustments in head position were made to match the DV coordinates. Virus was injected at AP –3.2, ML –0.9, and DV –4.4. A small hole was drilled into the skull and 230 nL of virus (see titer above) was injected through a pulled glass pipet using a Nanoject 2000 (Drummond Scientific; 10 pulses of 23 nL). At 5 minutes after injection, the glass pipet was slowly withdrawn over 5 min. After closing the skin with vicryl sutures, mice received 0.5 mL of 0.9% saline i.p. and were allowed to recover for >1 hour before being returned to their home cages. Animals were housed for at least 3 weeks after injection to allow AAV expression before being experiments.

### Antibodies and Reagents

| Name | Manufacturer | Catalog | RRID | Usage |
| --- | --- | --- | --- | --- |
| Mouse anti-V5 antibody | ThermoFisher | R960-25 | RRID:AB_2556564 | IHC, 1:1000<br>WB, 1: 1000 |
| Chicken anti-TH antibody | Millipore | AB9702 | RRID:AB_570923 | IHC, 1:500<br>ICC, 1:1000 |
| Rabbit anti-RFP antibody | Rockland | 600-401-379 | RRID:AB_2209751 | IHC, 1:500 |
| Goat anti-Chicken IgY (H+L), Alexa Fluor Plus 488 | ThermoFisher | A-32931TR | RRID:AB_2866499 | IHC/ICC,<br>1:1000 |
| Goat anti-Mouse IgG (H+L), Alexa Fluor 647 | ThermoFisher | A-21235 | RRID:AB_2535804 | IHC/ICC, 1:1000 |
| Goat anti-Rabbit IgG (H+L), Alexa Fluor 555 | ThermoFisher | A-21428 | RRID:AB_2535849 | IHC, 1:750 |
| Streptavidin-AlexaFluor647 | ThermoFisher | S-21374 | RRID:AB_2336066 | IHC, 1:400<br>ICC, 1:500 |
| Streptavidin HRP | Vector Laboratories | SA-5014 | RRID:AB_2336510 | WB, 3:10,000 |
| Goat anti-Mouse HRP | Jackson ImmunoResearch | 115-005-003 | RRID:AB_2338447 | WB, 1:5000 |
| Biotin tyramide (Biotin Phenol/BP) | Iris Biotech GmbH | LS3500 | NA | APEX2, 0.5 mM |
| Hydrogen Peroxide (H<sub>2</sub>O<sub>2</sub>) | MilliporeSigma | H1009 | NA | APEX2, 1 mM |

IHC: Staining of acute brain slices or sections, ICC: Staining of synaptosomes, WB: western blotting, APEX2: slice and synaptosome labeling concentrations

### Acute Brain Slice Preparation

Mice were anesthetized with euthasol and transcardially perfused with 10-15 ml of ice-cold cutting solution (92 mM NMDG, 2.5 mM KCl, 30 mM NaHCO_3_, 20 mM HEPES, 1.25 mM NaH_2_PO_4_, 2 mM thiourea, 5 mM sodium ascorbate, 3 mM sodium pyruvate, 10 mM MgSO_4_, 0.5 mM CaCl_2_, and 25 mM D-glucose) at pH 7.3-7.4 and saturated with 95% O_2_/5% CO_2_. Brains were rapidly extracted and placed into ice-cold cutting solution saturated with 95% O_2_/5% CO_2_. For coronal slices, the brain was split roughly in half with a coronal cut at the level of the mid-hypothalamus (approximately 1.5 mm posterior to Bregma), and the cut surfaces were glued directly to the vibratome stage (Leica VT1000S). For sagittal slices, brains were laterally split in half at the midline and the cut (medial) surfaces were glued to 2% agarose blocks angled at 11 degrees. 300 μm-thick slices were prepared in ice-cold cutting solution continuously saturated with 95% O_2_/5% CO_2_. The brainstem and cerebellum were removed from sagittal slices. A typical set of coronal or sagittal slices from a mouse are shown in **Figure 1 – figure supplement 2a**.

### Electrophysiological recordings

After preparation in cold cutting solution, slices were transferred to oxygenated normal ACSF containing 125.2 NaCl, 2.5 KCl, 26 NaHCO_3_, 1.3 MgCl_2_·6H_2_O, 2.4 CaCl_2_, 0.3 NaH_2_PO_4_, 0.3 KH_2_PO_4_, and 10 D-glucose (pH 7.4, 290 ± 5 mOsm) at 34 °C and allowed to recover for at least 40 min before the recordings. After recovery, slices were transferred to ACSF +/- 0.5 mM BP and incubated for 1 hr at room temperature. Electrophysiological recordings were performed on an upright Olympus BX50WI (Olympus, Tokyo, Japan) microscope equipped with a 40x water immersion objective, differential interference contrast (DIC) optics and an infrared video camera. Slices were transferred to a recording chamber and maintained under perfusion with normal ACSF (1.5-2 mL/min) at 34°C. All recorded dopaminergic neurons were located in the substantia nigra pars compacta and identified by larger somatic size than neighboring neurons. Patch pipettes (3-5 MΩ) were pulled using P-97 puller (Sutter instruments, Novato, CA) and filled with internal solutions contained (in mM): 115 K-gluconate, 10 HEPES, 2 MgCl_2_, 20 KCl, 2 MgATP, 1 Na_2_-ATP, and 0.3 GTP (pH = 7.3; 280 ± 5 mOsm).

Cell-attached patch clamp recordings were performed with a MultiClamp 700B amplifier (Molecular Devices, Forster City, CA) and digitized at 10 kHz with InstruTECH ITC-18 (HEKA, Holliston, MA). Once obtaining stable patch configuration, spontaneous firing was recorded for 3 min. Data was acquired using WINWCP software (developed by John Dempster, University of Strathclyde, UK) and analyzed using Clampfit (Molecular Devices), and Igor Pro (Wavemetrics, Lake Oswego, OR).

### APEX2 biotinylation in brain slices

After preparation in cold cutting solution, slices were transferred to jars containing 70 mL of aCSF (125.2 mM NaCl, 2.5 mM KCl, 26 mM NaHCO_3_, 1.3 mM MgCl_2_·6H_2_O, 2.4 mM CaCl_2_, 0.3 mM NaH_2_PO_4_, 0.3 mM KH_2_PO_4_, 10 mM D-Glucose) supplemented with 0.5 mM biotin-phenol and 1 μM tetrodotoxin. aCSF was continuously saturated with 95% O_2_/5% CO_2_. Slices were allowed to recover for 60 minutes at room temperature. After recovery, APEX2 labeling was initiated by the addition of 1 mM H_2_O_2_ to aCSF at room temperature. We tested labeling periods of 1-5 minutes (**Figure 1 – figure supplement 2b-d**) and found 3 minutes to be sufficient for downstream applications. To quench labeling, slices were rapidly transferred to a separate jar containing 50 mL of quenching aCSF (aCSF supplemented with 10 mM Trolox, 20 mM sodium ascorbate, and 10 mM NaN_3_). After 5 minutes rest at room temperature, slices were either transferred to fixative solution for downstream immunostaining, or transferred to ice-cold quenching aCSF for rapid dissection and downstream western blotting/proteomics.

### Synaptosome Preparation and APEX2 biotinylation

Synaptosomes were prepared using standard procedures (Gray & Whittaker, 1962). Mice were sacrificed by cervical dislocation, after which forebrains were rapidly dissected and placed in 10 volumes of ice-cold buffer consisting of 0.32 M sucrose, 4 mM HEPES pH 7.4, and protease inhibitors (complete^™^ EDTA-free protease inhibitor, Roche). Tissue was homogenized on ice in a glass-glass dounce homogenizer with 10 gentle strokes of loose and tight clearance pestles. All subsequent purification steps were performed on ice or at 4°C unless otherwise specified. The homogenate was centrifuged at 1,000 xg (Eppendorf 5424R) for 10 min to remove nuclei and cellular debris, yielding a P1 pellet and an S1 supernatant. The S1 supernatant was further centrifuged at 10,000 x g for 15 min to obtain the crude synaptosome pellet (P2). The P2 pellet was resuspended in sucrose buffer, incubated for 5 minutes on ice, and re-centrifuged at 10,000 x g for 15 min. The washed P2 pellet was resuspended in phosphate-buffered saline (PBS) with 0.5 mM BP and incubated for 30 minutes at room temperature. For immunostaining experiments, the 30-minute incubation in PBS + BP occurred on poly-lysine coated coverslips to allow synaptosome sedimentation and adherence. Unbound synaptosomes were removed by several brief washes in PBS + BP prior to APEX2 labeling. APEX2 labeling was initiated by the addition of 0.5x volumes of 2 mM H_2_O_2_ (1 mM final). After 60 seconds, labeling was quenched by addition 4x volumes of ice-cold PBS + 12.5 mM Trolox, 25 mM sodium ascorbate, and 12.5 mM NaN_3_ (10, 20, and 10 mM final, respectively). For immunostaining, synaptosomes were washed several more times in quenching PBS over 5 minutes, followed by addition of fixative solution. For western blotting and proteomics, synaptosomes were collected by re-centrifugation at 10,000 x g for 15 min, flash frozen on liquid nitrogen, and stored at −80°C until further use.

### Histology and Immunofluorescence

For initial characterization of AAV5-CAG-DIO-APEX2NES specificity, mice were anesthetized with euthasol and transcardially perfused with ~15 mL of 0.9% saline followed by 40-50 mL of ice-cold 4% paraformaldehyde (PFA) in 0.1 M phosphate buffer (PB), pH 7.4. Brains were post-fixed in 4% PFA in 0.1 M PB for 6-12 hours at 4°C, washed three times in phosphate buffered saline (PBS), and sectioned at 50 μm on a Leica VT1000S vibratome. Sections were placed in cryoprotectant solution (30% ethylene glycol, 30% glycerol, 0.1 M PB, pH7.4) and stored at −20°C until further use.

Sections were removed from cryoprotectant solution and washed three times in tris-buffered saline (TBS) at room temperature. Sections were then permeabilized in TBS + 0.3% Triton-X 100 for one hour at room temperature, followed by blocking in TBS + 10% normal goat serum (NGS) and 0.3% Triton-X 100 for 1.5 hours at room temperature. Sections were then directly transferred to a pre-chilled solution containing primary antibodies in TBS + 2% NGS + 0.1% Triton-X 100 and incubated overnight at 4°C. Sections were washed in TBS five times over an hour at room temperature. Sections were incubated in a solution containing secondary antibodies in TBS + 2% NGS + 0.1% Triton-X 100 at room temperature for 1.5 hours, followed by four washes in TBS+T over 45 minutes at room temperature. Following four additional washes in TBS, sections were slide mounted and coverslipped with Fluoromount G (Southern Biotech). See *Antibodies and Reagents* for a complete list of antibodies and concentrations used in this study.

For immunostaining of acute brain slices after APEX2 labeling, slices were transferred to ice-cold 4% paraformaldehyde in 0.1M phosphate-buffer + 4% sucrose and fixed overnight at 4°C. To remove lipids and enhance antibody penetration, fixed slices were transferred to CUBIC solution 1A, consisting of 10% wt Triton X-100, 5% wt NNNN-tetrakis (2-HP) ethylenediamine, 10% wt Urea, and 25 mM NaCl (Susaki & Ueda, 2016). Slices were blocked in TBS + 10% normal goat serum (NGS) and 0.3% Triton-X 100 for XX hour at room temperature and then transferred to TBS + 2% NGS and 0.3% Triton-X 100 supplemented with primary antibodies. After 72 hours incubation at 4°C with primary antibodies, slices were washed five times over 10 hours in TBS. Slices were incubated for 24 hours at room temperature in TBS + 2% NGS + 0.3% Triton-X 100 supplemented with secondary antibodies and fluorophore-conjugated streptavidin. After five washes in TBS over 10 hours, sections were slide mounted and coverslipped with Fluoromount G. See *Antibodies and Reagents* for a complete list of antibodies and concentrations used in this study.

For immunostaining of synaptosomes after APEX2 labeling, synaptosomes adhered to poly-lysine coverslips were fixed with 4% paraformaldehyde in 0.1 M phosphate-buffer + 4% sucrose for 10 minutes at room temperature. After several washes in PBS, synaptosomes were incubated with PBS + 0.1M glycine + 0.05% Tween-20 for 15 minutes at room temperature. After blocking/permeabilization with PBS + 10% NGS + 0.2% Tween-20 for one hour at room temperature, synaptosomes were incubated with primary antibodies in PBS + 2% NGS + 0.1% Tween-20 at 4°C overnight. After three washes in PBS, synaptosomes were incubated in secondary antibodies in PBS + 2% NGS + 0.1% Tween-20 for 1 hour at room temperature. After three more washes in PBS, coverslips were stored and imaged in Fluoromount G. See *Antibodies and Reagents* for a complete list of antibodies and concentrations used in this study.

### Tissue lysis and protein processing

Capture and processing of biotinylated proteins was conducted as previously described (Kalocsay, 2019; G. Liu et al., 2020) with only minor modifications. Immediately after dissection in ice-cold quenching aCSF, tissues were flash frozen in liquid nitrogen and stored at −80°C until further use. Frozen tissues or synaptosome pellets were homogenized on ice in a glass dounce homogenizer (Sigma D9063) with 30 strokes of both A and B pestles. Lysis was in 0.75 mL of ice-cold tissue lysis buffer, consisting of 50 mM Tris pH 8.0, 150 mM NaCl, 10 mM EDTA, 1% Triton X-100, 5 mM Trolox, 10 mM sodium ascorbate, 10 mM sodium azide, and 1x EDTA-free protease inhibitors (Roche). After addition of 39 μL of 10% SDS (final concentration 0.5%), lysates were rotated for 15 minutes at 4°C. Lysates were clarified by centrifugation at 21,130 x g for 10 minutes at 4°C. Supernatants were transferred to a new pre-chilled Eppendorf tube for trichloroacetic acid (TCA) precipitation (for mass spectrometry) or stored at −80°C (for western blotting).

Proteins were precipitated from lysates by the addition of an equal volume of ice-cold 55% TCA. Samples were incubated on ice for 15 minutes, followed by centrifugation at 21,130 x g for 10 minutes at 4°C. Protein pellets were resuspended in 1 mL of acetone pre-chilled to −20°C and re-centrifuged as before. Pellets were resuspended and re-centrifuged another three times in 1 mL of acetone pre-chilled to −20°C, for a total of four washes. Residual acetone was removed, and protein pellets were resuspended in Urea Dissolve Buffer (8M Urea, 1% SDS, 100 mM sodium phosphate pH 8, 100 mM NH_4_HCO_3_). Dissolution of pellets was facilitated by water both sonication for 10 minutes followed by gentle agitation on an orbital shaker for 1 hour at room temperature. Removal of residual TCA was confirmed by checking that the pH ~8.0. In some cases, a small aliquot (5%) of the resuspended protein was flash-frozen and stored at −80°C. 1/49 the volume of 500 mM TCEP (Sigma, cat. #646547) and 1/19 the volume of freshly prepared 400 mM iodoacetamide (ThermoFisher, cat. #90034) in 50 mM NH_4_HCO_3_ was added to the protein resuspension for disulfide reduction and cysteine alkylation at final concentrations of 10 mM TCEP and 20 mM iodoacetamide. The suspension was vortexed and incubated in the dark for 25 minutes at room temperature. Alkylation was quenched by addition of 1/19 the volume of 1 M DTT to reach 50 mM DTT. Samples were diluted with 0.87x volumes of H_2_O to reach a final concentration of 4 M urea and 0.5% SDS.

### Capture of biotinylated proteins for mass spectrometry

Streptavidin magnetic beads (ThermoFisher #88817) were resuspended and washed three times in Urea Detergent Wash Buffer (4 M Urea, 0.5% SDS, 100 mM sodium phosphate pH 8) for at least 10 minutes at 4°C. After washing, streptavidin beads were resuspended in ice-cold Urea Detergent Wash Buffer and 50 μL containing 0.5 mg of beads was added to each sample. Proteins were incubated with streptavidin beads overnight on a rotor at 4°C. After 14-18 hours, the unbound supernatant was discarded, and beads were resuspended in 1 mL of Urea Detergent Wash Buffer and transferred to a new tube. Beads were washed three times for 5-10 minutes in 1 mL of Urea Detergent Wash Buffer at room temperature. After the third wash, beads were resuspended in 1 mL of Urea Wash Buffer (4 M Urea, 100 mM sodium phosphate pH 8) and transferred to a new tube. After three 5-10-minute washes in 1 mL of Urea Wash Buffer at room temperature, beads were resuspended in 200 μL of Urea Wash Buffer and transferred to a new tube. A 10 μL aliquot (5%) was transferred to a separate tube for western blotting, and the remaining 190 μL of buffer were removed on a magnetic stand. Beads were flash frozen and stored at −80°C.

### Western blotting

The protein concentration of frozen tissue lysates was determined using the BCA assay (Pierce, ThermoFisher catalog #23225) and diluted with 4x LDS sample buffer (ThermoFisher, catalog #NP0007) supplemented with 20 mM DTT and boiled for 5 min at 95°C. Frozen streptavidin beads were resuspended in ~ 20 μL of 1x LDS sample buffer supplemented with 20 mM DTT and 2 mM biotin. Samples were boiled for 5 min at 95°C to elute biotinylated proteins. Beads were immediately placed immediately onto a magnetic rack and the entire sample was immediately loaded into 10% Bis-Tris polyacrylamide gels (Invitrogen, ThermoFisher catalog #NP0303BOX) and transferred to PVDF membranes (Immobilon-P, MilliporeSigma, catalog #IPVH00010). Membranes were initially washed for 15 minutes in TBST (1X TBS + 0.1% Tween 20), blocked for an hour in 5% BSA/TBST, and incubated overnight at 4°C with primary antibody in 5% bovine serum albumin/TBST overnight. After primary incubation, membranes were washed three times in TBST prior to incubation with streptavidin-HRP or HRP-conjugated secondary antibody in 2.5% BSA/TBST for one hour at room temperature. After secondary incubation, membranes were washed three times in TBST. Signal was developed using Immobilon enhanced chemiluminescent substrate (Millipore, catalog #WBKLS0500) and imaged on an Azure Biosystems C600 system.

### On bead digestion and liquid chromatography/tandem mass spectrometry (LC-MS/MS)

Proteins bounded streptavidin beads were resuspended in 200 μl of digestion buffer (1M urea, 100mM EPPS pH 8.5,4% acetonitrile) and digested with 2 μg of trypsin/LysC mix overnight at 37°C. The next day, digested peptides were collected in a new microfuge tube and digestion was stopped by the addition of 1% TFA (final v/v), followed by centrifugation at 14,000 x g for 10 min at room temperature. Cleared digested peptides were desalted on a SDB-RP Stage-Tip and dried in a speed-vac. Dried peptides were dissolved in 3% acetonitrile/0.1% formic acid. Desalted peptides (300-500 ng) were injected onto an EASY-Spray PepMap RSLC C18 50 cm x 75 μm column (Thermo Scientific), which was coupled to the Orbitrap Fusion Tribrid mass spectrometer (Thermo Scientific). Peptides were eluted with a non-linear 120 min gradient of 5-30% buffer B (0.1% (v/v) formic acid, 100% acetonitrile) at a flow rate of 250 nL/min. The column temperature was maintained at a constant 50 °C during all experiments.

Samples were run on the Orbitrap Fusion Tribrid mass spectrometer with a data independent acquisition (DIA) method for peptide MS/MS analysis (Bruderer et al., 2015). Survey scans of peptide precursors were performed from 350-1200 *m/z* at 120K FWHM resolution (at 200 *m/z)* with a 1 x 10^6^ ion count target and a maximum injection time of 60 ms. After a survey scan, 26 m/z DIA segments acquired at from 200-2000 *m/z* at 60K FWHM resolution (at 200 *m/z)* with a 1 x 10^6^ ion count target and a maximum injection time of 118 ms. HCD fragmentation was applied with 27% collision energy and resulting fragments were detected using the rapid scan rate in the Orbitrap. The spectra were recorded in profile mode.

### Raw mass spectrometry data processing

DIA data were analyzed with directDIA 2.0, a spectral library-free analysis pipeline featured in Spectronaut Pulsar X software (Biognosys AG). The default settings were used for targeted analysis of DIA data in Spectronaut except the decoy generation was set to “mutated”. False discovery rate (FDR) was estimated using the mProphet approach (Reiter et al., 2011) and set to 1% at peptide precursor level and at 1% at protein level. For peptides and proteins that were not detected in a given sample (directDIA output as ‘Filtered’), the intensity was set to 0 for downstream analysis.

### Proteomic Differential Expression Analysis and Filtering

Total intensity normalized protein abundances were used for all differential expression analyses, which consisted of a Welch’s (unequal variance) t-test with Benjamini-Hochberg procedure to control the False discovery rate (FDR). For visualization and clustering, total intensity normalized protein abundances were log2 transformed after adding 1.

A graphical summary of the initial filtering for VM and striatum APEX2 proteomics data is shown in **Figure 3 – figure supplement 1d**. Most filters were based on the APEX2 proteomics data alone, with some additional filters based on DA neuron scRNA-seq data derived from Saunders et al. (2018) (see below). VM proteins were filtered as follows:

1. Proteins meeting statistically significant (FDR < 0.05) enrichment in APEX2^+^ vs. APEX2^-^ (Control) differential expression were retained
2. Proteins were retained if they met either one of the two following conditions (a OR b):

a. Mean DA neuron scRNA-seq expression above the lower bound (mean – standard deviation)
b. Statistically significant (FDR < 0.05) enrichment in APEX2^+^ vs. APEX2^-^ (Control) differential expression in all three regions (VM, MFB, and Str)
3. Proteins were removed if they were encoded by genes with very low DA neuron-specificity in scRNA-seq data (Mann-Whitney U-test comparing DA neurons vs. all other midbrain cells).

Striatum proteins were filtered as follows:

1. Proteins meeting statistically significant (FDR < 0.05) enrichment in APEX2^+^ vs. APEX2^-^ (Control) differential expression were retained
2. Proteins were retained if they met any of the following conditions (a OR b OR c):

a. Statistically significant enrichment (FDR < 0.05) in APEX2^+^ vs. APEX2^-^ (Control) differential expression for either VM or MFB
b. log2 Fold Change > 1 in APEX2^+^ vs. APEX2^-^ (Control) comparisons for both VM and MFB samples
c. High mean DA neuron scRNA-seq expression (above the mean plus standard deviation) and DA neuron specificity (Mann-Whitney U-test comparing DA neurons vs. all other midbrain cells or vs. all striatal cells)
3. Proteins were retained if they met either one of the two following conditions (a OR b):

a. Mean DA neuron scRNA-seq expression above the lower bound (mean minus standard deviation)
b. Statistically significant (FDR < 0.05) enrichment in APEX2^+^ vs. APEX2^-^ (Control) differential expression in all three regions (VM, MFB, and Str)
4. Proteins were removed if they were encoded by genes with very low DA neuron-specificity in scRNA-seq data (Mann-Whitney U-test comparing DA neurons vs. all other midbrain cells or vs. all striatal cells).

### Gene Ontology (GO) Analysis

For all gene ontology analyses, a single list of unique genes encoding the corresponding proteins was used (i.e., a single gene entry was used when multiple protein isoforms in the list were encoded by a single gene). For peptides/protein groups mapped to multiple, homologous proteins, the first gene entry as determined by Spectronaut default settings was used. The VM vs. striatum GO analysis shown in **Figure 3d** was conducted using web-based Enrichr (Xie et al., 2021) with 2018 GO Terms for Cellular Component, Biological Process, and Molecular Function (Ashburner et al., 2000; Gene Ontology Consortium, 2021). Enrichr was also used for subcellular compartments analysis shown in **Figure 3 – figure supplement 3a-b**. Additional targeted gene ontology analysis shown in **Figure 3 – figure supplement 3c-d** was conducted manually using the *SciPy* implementation of the hypergeometric test. Nuclear-related ontologies were obtained from COMPARTMENTS (Binder et al., 2014) and mitochondrial localization ontologies were obtained from MitoCarta 3.0 (Rath et al., 2021). The synaptic gene ontology analysis shown in **Figure 4** and **Figure 4 – figure supplement 1** was conducted using SynGO (Koopmans et al., 2019).

### Image acquisition and analysis

Imaging of 50 μm sections from perfusion-fixed brain was conducted on a Nikon Ti2 Eclipse epifluorescence microscope or on a Leica SP8 scanning confocal microscope. To confirm the specificity of V5 expression in DA neurons, tile scan epifluorescence images of the entire ventral midbrain were collected at 2-3 z-planes per section. V5-positive neurons were first identified using only the V5 channel and their somas were segmented as ROIs. Each V5-positive neuronal ROI was subsequently scored for tdTomato and TH expression. Neurons within both the substantia nigra pars compacta (SNc) and ventral tegmental area (VTA) were quantified. High resolution (60x/1.4 NA) confocal images of striatal sections confirmed that V5 (APEX2), tdTomato, and TH were localized exclusively within dopaminergic axons. 300 μm thick brain slices were imaged only by confocal microscopy. To assess biotin labeling throughout the slice, Z-stacks spanning the entire slice depth were acquired at 4.18 μm intervals using a 20x / 0.4 NA objective.

### scRNA-seq Analysis

For comparison to scRNA-seq, we obtained Drop-seq count matrices for substantia nigra and striatum from GSE116470 (DropViz, Saunders et al., 2018). To identify dopamine neurons, we first performed unsupervised clustering on the substantia nigra count matrices using the Phenograph (Levine et al., 2015) implementation of Louvain community detection after selection of highly variable genes and construction of a k-nearest neighbors graph as described previously (Levitin et al., 2019). We identified a single cluster with statistically significant co-enrichment of dopamine neuron markers such as *Th* and *Slc6a3* based on the binomial test for expression specificity (Shekhar et al., 2016) as shown in the UMAP (Becht et al., 2018) embedding in **Figure 5 – figure supplement 1a**. After sub-clustering the putative dopamine neurons using the methods described above, we identified a small sub-cluster with statistical enrichment of astrocyte markers such as *Agt*, *Gja1, Glul*, and *Slc1a3*. We discarded this sub-cluster as likely astrocyte contamination and removed all remaining cells with fewer than 1,000 unique transcripts detected to produce a count matrix of high-confidence dopamine neuron profiles. We sub-clustered these profiles to identify five transcriptionally distinct dopamine neuron subsets with markers determined using the binomial test shown in **Figure 5 – figure supplement 1b**.

We used these high-confidence profiles to identify genes with enrichment in dopamine neurons compared to all midbrain cells by differential expression analysis as described in Szabo, Levitin et al., (2019) with minor modifications. Briefly, to perform differential expression analysis between two groups of cells, we randomly sub-sampled the data so that both groups are represented by the same number of cells. Next, we randomly sub-sampled the detected transcripts so that both groups have the same average number of transcripts per cell. Finally, we normalized the two sub-sampled count matrices using *scran* (Lun et al., 2016) and analyzed differential expression for each gene using the SciPy implementation of the Mann-Whitney U-test. We corrected the resulting p-values for false discovery using the Benjamini-Hochberg procedure as implemented in the *statsmodels* package in Python. We used differential expression analyses between high-confidence dopamine neurons and the remaining cells in the midbrain to select genes with >8-fold specificity for expression in DA neurons (log2FC > 3 and FDR < 0.01).

### Ana/ysis of Proteins Encoded by DA Neuron Marker Genes and PD-linked Genes

DA neuron marker genes (genes specific to mouse dopamine neurons) were identified using scRNA-seq data as described above. Out of 64 genes with >8-fold specificity for expression in DA neurons, 55 corresponding proteins were present in the filtered APEX2 proteomics data. See **Figure 5 – source data 1** for complete summary of mouse DA neuronal marker genes, corresponding mouse proteins, and protein abbreviations shown in **Figure 5b**. Human genes linked to PD via monogenic inheritance were obtained and classified according to the guidelines put forth in Marras et al. (2016). Human genes linked to PD via GWAS were obtained from the most recent meta-analysis (Nalls et al., 2019). The closest gene to each SNP in Table S2 was used for downstream analysis. For all human genes, mouse gene orthologs were obtained via the web-based ID conversion tool (BioMart, Ensembl). See **Figure 5 – source data 2** for complete summary of human PD-linked genes, mouse gene orthologs, mouse proteins, and protein abbreviations shown in **Figure 5c**.

### Visualization and Statistical Analysis

Cartoon graphics (e.g., **Figure 3d** and **Figure 4f**) were created in Adobe Illustrator 24.3 (Adobe, Inc.) with additional illustrations from BioRender (https://biorender.com/). Proteins were selected for display on this basis of inclusion in significantly over-represented GO Terms, with additional proteins selected based on manual curation of the relevant literature. Unless otherwise noted, all proteins displayed were present in the filtered APEX2 proteomics data. Protein abbreviations and corresponding full protein names are provided as source data for each respective figure.

Unless otherwise noted, all statistical analysis and data visualization was conducted in Python using *SciPy, Matp/ot/ib*, and *Seaborn* packages. For visualization of proteomics data (log-log abundance plots, Z-scores, clustered heatmaps, etc.), total intensity normalized protein abundances were log2 transformed after adding 1. The total intensity normalized, log2 transformed protein intensities are generally referred to as log2 protein abundance, as specified in figure captions. For clustered heatmaps, Z-scores of log2 protein abundances were first calculated using the *zscore* function within the *SciPy Stats* module, after which the row and column clustering was calculated using the *linkage* function (metric = ‘Euclidean’, method = ‘average’) within *fastcluster 1.2.3* (Müllner, 2013) and passed to *Seaborn clustermap*.

## Materials Availability

There are restrictions to the availability of AAV5-CAG-DIO-APEX2NES virus due to limited production size. The exact plasmid used for production of this virus (AAV-CAG-DIO-APEX2NES, Addgene plasmid #79907) can be ordered from Addgene and/or sent directly to Vector BioLabs for further production.

## Data Availability

The mass spectrometry proteomics data have been deposited to the ProteomeXchange Consortium via the PRIDE (Perez-Riverol et al., 2019) partner repository with the dataset identifier PXD026229. Raw label-free quantification intensity values for proteomics data can be found in **Figure 2 — source data 2**. The scRNA-seq data analyzed are publicly available as GSE116470 (Saunders et al., 2018). High confidence DA neuron profiles used in this study are reported in **Figure 5 — source data 3**.

## Code Availability

Python code used for clustering and visualization of scRNA-seq data can be found at www.github.com/simslab/cluster_diffex2018 and for differential protein expression analysis of mass spectrometry data at www.github.com/simslab/proteomics2021.

## Competing interests

The authors declare that no competing interests exist.

## Contributions

BDH conceived the overall project and APEX2 slice labeling procedures. BDH executed all mouse APEX2 experiments, with assistance from SJC in transcardial perfusion and brain slice preparation. SJC performed electrophysiological recordings. BDH executed protein purification, immunoprecipitation, western blotting, and immunofluorescence experiments. RKS executed all on-bead digestion and mass spectrometry experiments. PAS conducted scRNA-seq analysis and visualization. BDH conducted proteomics data analysis with input from PAS. PAS and DS supervised the research. BDH wrote the manuscript with input from PAS and DS. All authors edited, read, and approved the final manuscript.

## Acknowledgements

This work was conducted in collaboration with the Proteomics Shared Resource within the Herbert Irving Comprehensive Cancer Center at Columbia University Irving Medical Center (NIH Grant 2P30 CA013696-45). This research was funded in part by Aligning Science Across Parkinson’s [ASAP-000375] (DS and PAS) through the Michael J. Fox Foundation for Parkinson’s Research (MJFF). For the purpose of open access, the author has applied a CC BY public copyright license to all Author Accepted Manuscripts arising from this submission. This work was supported by the JPB Foundation (DS). This work was supported by NIH grants F30 DA047775-03 (BDH), R01 NS0954 (DS), R01 DA07418 (DS), and R01 MH122470 (DS). We would like to thank Eugene Mosharov for insightful discussions and Vanessa Morales for assistance with animal colony management.

## Notes

https://www.ebi.ac.uk/pride/archive/projects/PXD026229

### Competing Interest Statement

The authors have declared no competing interest.

### Summary of Updates

The manuscript has been revised with updated references to the literature and supportive electrophysiological recording data.

