## Supplementary Figures for "Subcellular proteomics of dopamine neurons in the mouse brain reveals axonal enrichment of proteins encoded by Parkinson’s disease-linked genes"

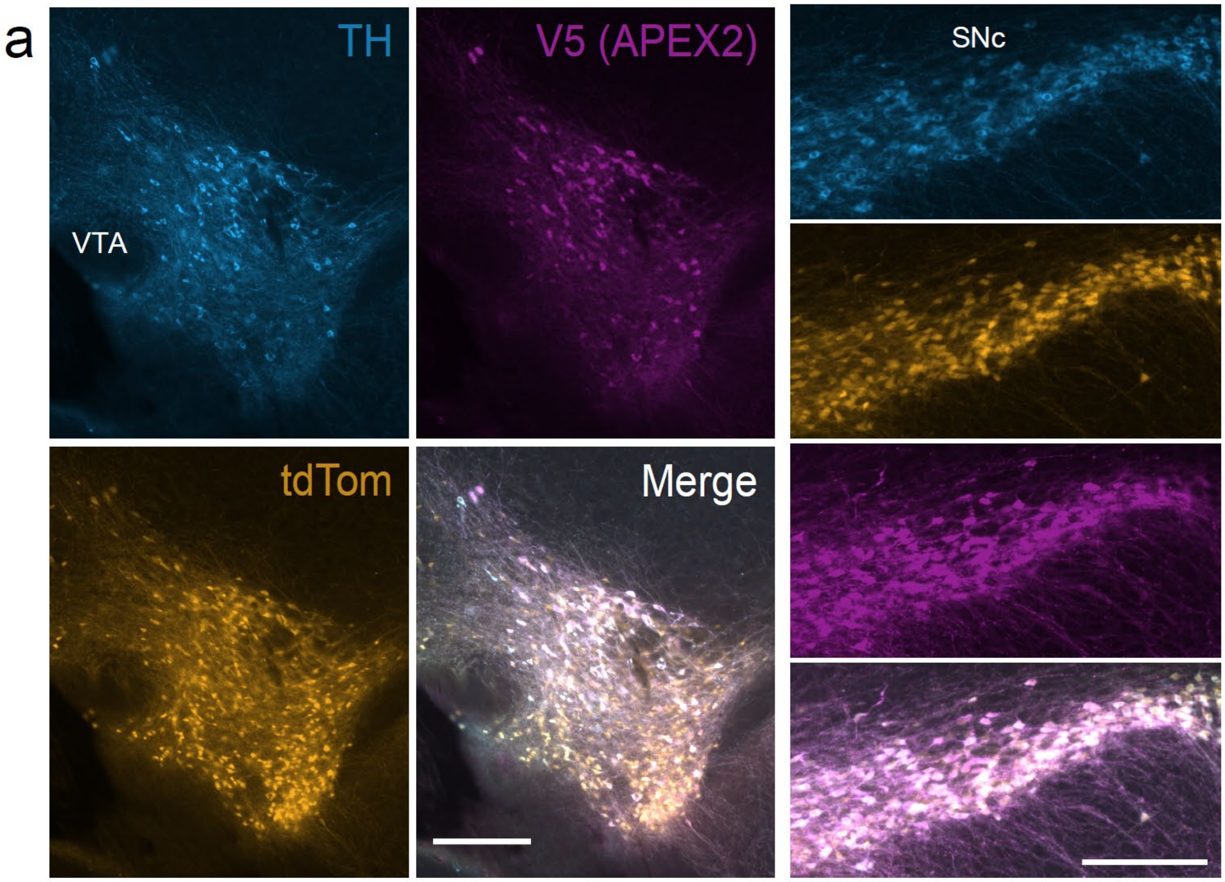


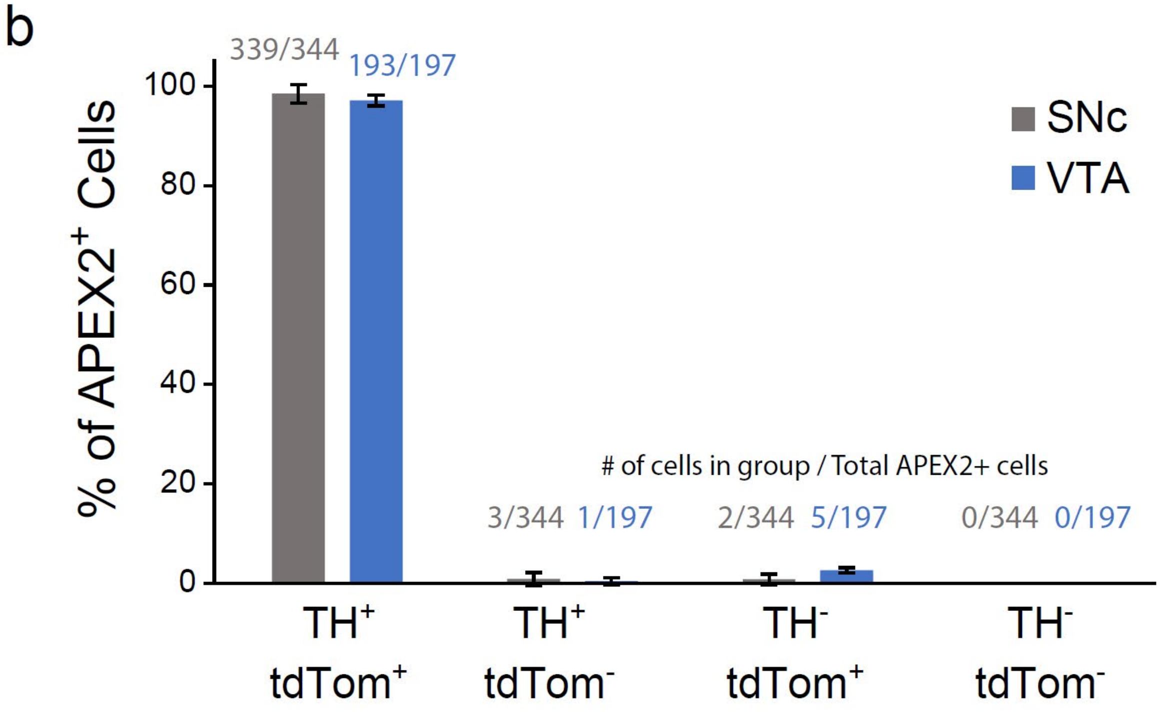


**Figure 1 – figure supplement 1**: **Specificity of APEX2-NES AAV expression**

(**a**) Immunostaining of mDA neurons in DAT$^{IRES\text{-}Cre}$/Ai9$^{tdTomato}$ mice injected with AAV5-CAG-DIO-APEX2-NES. The vast majority of V5-APEX2$^{+}$ neurons are TH$^{+}$/tdTomato$^{+}$. *Left*, ventral tegmental area, *right*: substantia nigra pars compacta, both scale bars: 250 µm.

(**b**) Quantification and cell counts related to (**a**), data from 2 sections each from n = 3 mice.

*Abbreviations*: (TH) tyrosine hydroxylase, (SNc) substantia nigra, (VTA) ventral tegmental area.

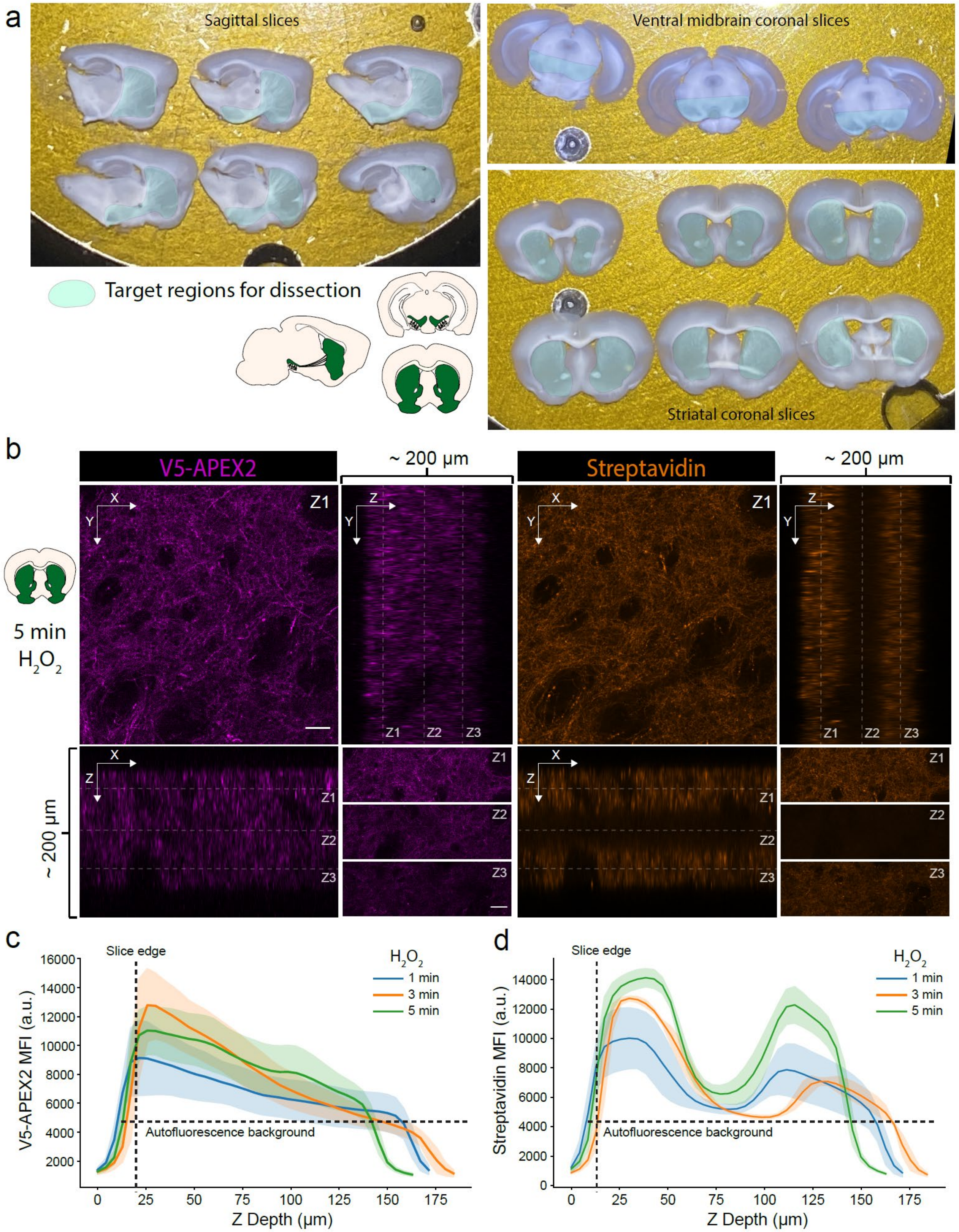


**Figure 1 – figure supplement 2**: **Characterization of slice labeling**

(**a**) Typical sets of sagittal or coronal slices. Target regions for anatomical dissection of mDA neuronal compartments are indicated.

(**b**) Anti-V5 (APEX2) and streptavidin-AlexaFluor647 staining of a cleared striatal slice after 5 minutes labeling with 1 mM hydrogen peroxide. *Upper left,* images display XY view at Z1 plane as indicated in the ZX (*lower* *left*) and YZ (upper right) views, scale bar: 50 µm. *Lower right*, top, middle, and bottom third of XY view at Z1, Z2, and Z3 planes, respectively, as indicated in the ZX and YZ views, scale bar: 50 µm.

(**c-d**) Anti-V5 (APEX2) and streptavidin-AlexaFluor647 mean fluorescence intensity as a function of Z depth and hydrogen peroxide ($H_2O_2$) labeling time. Mean ± standard deviation is plotted from 3 fields. Although V5-APEX2 fluorescence decreases throughout the slice depth (likely due to laser power attenuation), streptavidin fluorescence exhibits a bimodal distribution with sparse labeling in the center third.

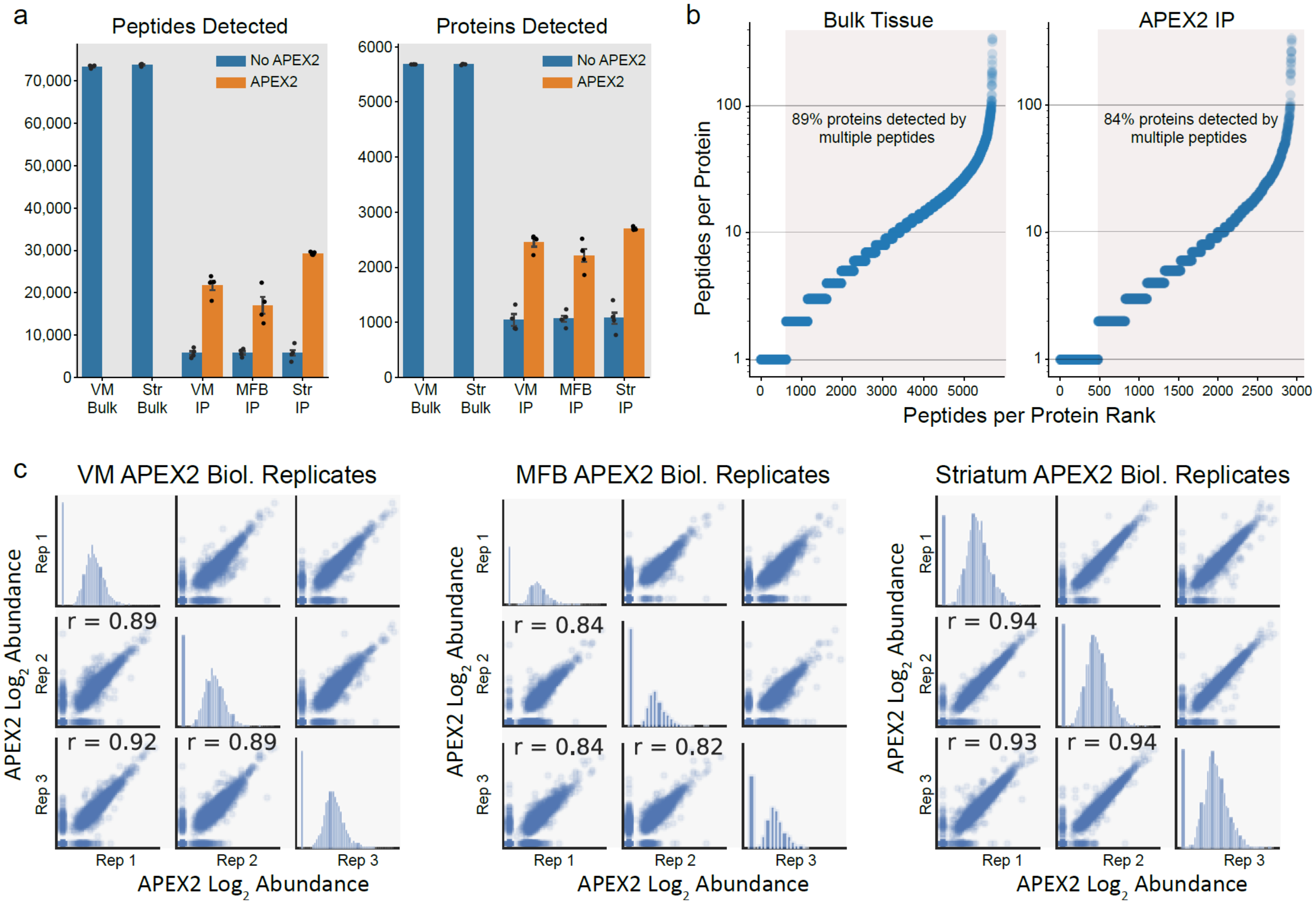


**Figure 2 – figure supplement 1**: **Proteomic depth and reproducibility**

(**a**) Mean ± SEM  of peptides and proteins detected per biological replicate of each sample type (n = 4 each). Tissue regions and protein fractions (bulk tissue or streptavidin IP) are indicated.

(**b**) Peptide coverage of detected proteins in each sample type (bulk tissue or streptavidin IP). Highlighted region shows proteins with >1 peptide.

(**c**) Pearson correlation (r) of three biological replicates ($APEX2^+$ streptavidin IP) samples from the indicated regions. Abundance represents $\log_2$(normalized intensity + 1) for each sample.

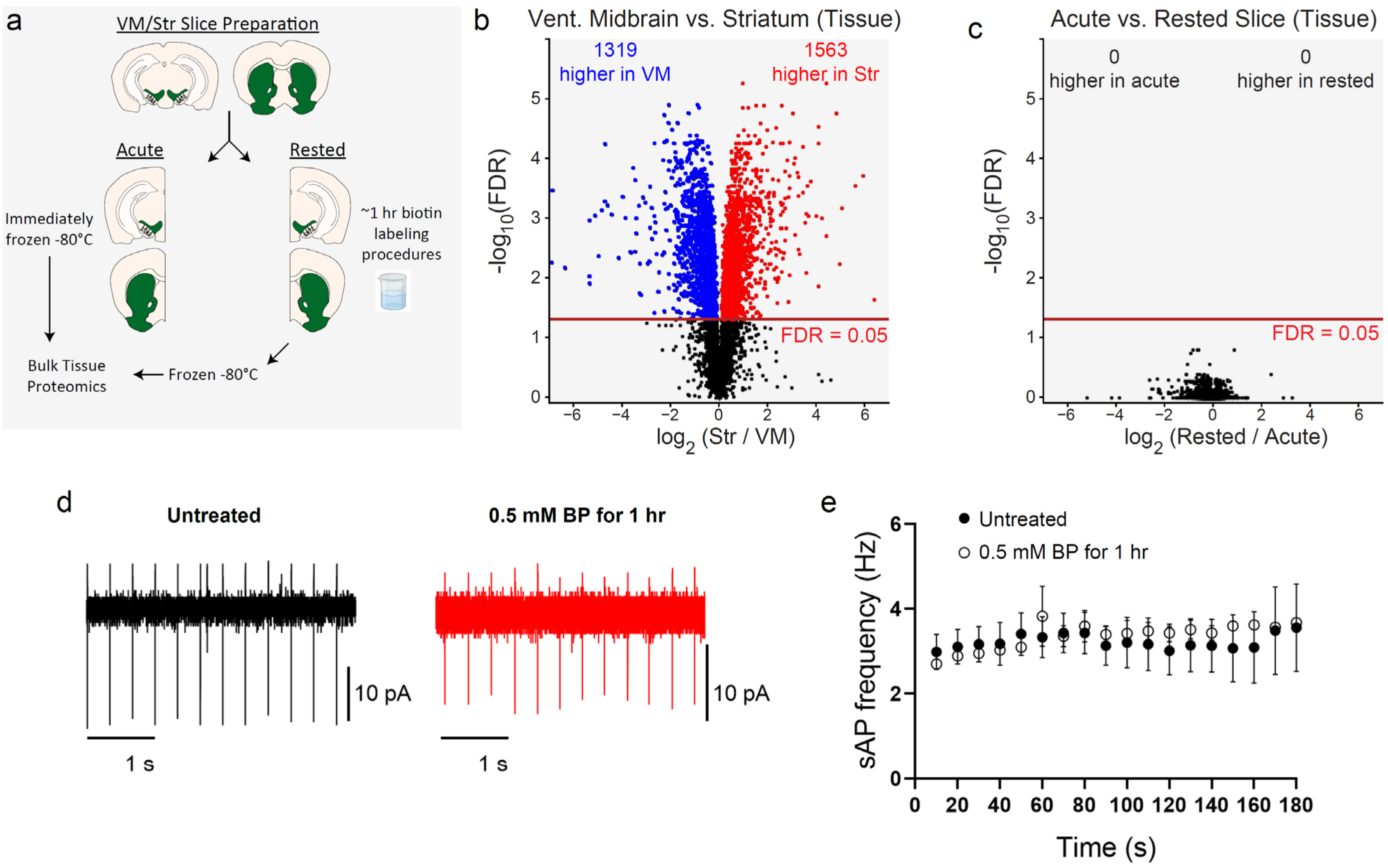


**Figure 2 – figure supplement 2: APEX2 slice labeling conditions do not distort bulk tissue proteomes or disrupt spontaneous firing in dopamine neurons**

(**a**) Schematic depicting bulk tissue proteomics experimental design. Coronal slice prepared from the midbrain or striatum were split into lateral halves: one half was immediately frozen (acute), while the other half was subjected to APEX2 labeling conditions (1-hour aCSF + 0.5 mM biotin-phenol followed by 3 minutes of 1 mM $H_2O_2$) and then frozen. See **Figure 2 – source data 2** for raw label-free quantification intensity values of peptides and proteins for all samples used in this study.

(**b**) Volcano plot comparing striatum and ventral midbrain bulk tissue proteomes. False discovery rate (FDR) represents q values from Benjamini-Hochberg procedure on Welch's (unequal variance) t-test. See **Figure 2 – source data 3** for complete results.

(**c**) Volcano plot comparing acute vs. rested bulk tissue proteomes for both striatum and ventral midbrain. False discovery rate (FDR) represents q values from Benjamini-Hochberg procedure on Welch's (unequal variance) t-test. See **Figure 2 – source data 3** for complete results.

(**d**) Cell attached recording of dopaminergic neurons in the substantial nigra. Representative traces of spontaneous firing of dopaminergic neurons from BP-treated and untreated slices.

(**e**) 1 hour incubation with 0.5 mM BP does not affect the spontaneous action potential firing (Untreated group: n=7 cells, N=2 mice, BP-treated: n=4 cells, N=2 mice. No significant effects were observed in two-way ANOVA: (Time) $F_{(17,100)} = 0.218$, p = 0.99, (Treatment) $F_{(1,100)} = 0.589$, p = 0.44, (Time:Treatment) $F_{(17,100)} = 0.098$, p = 0.99.

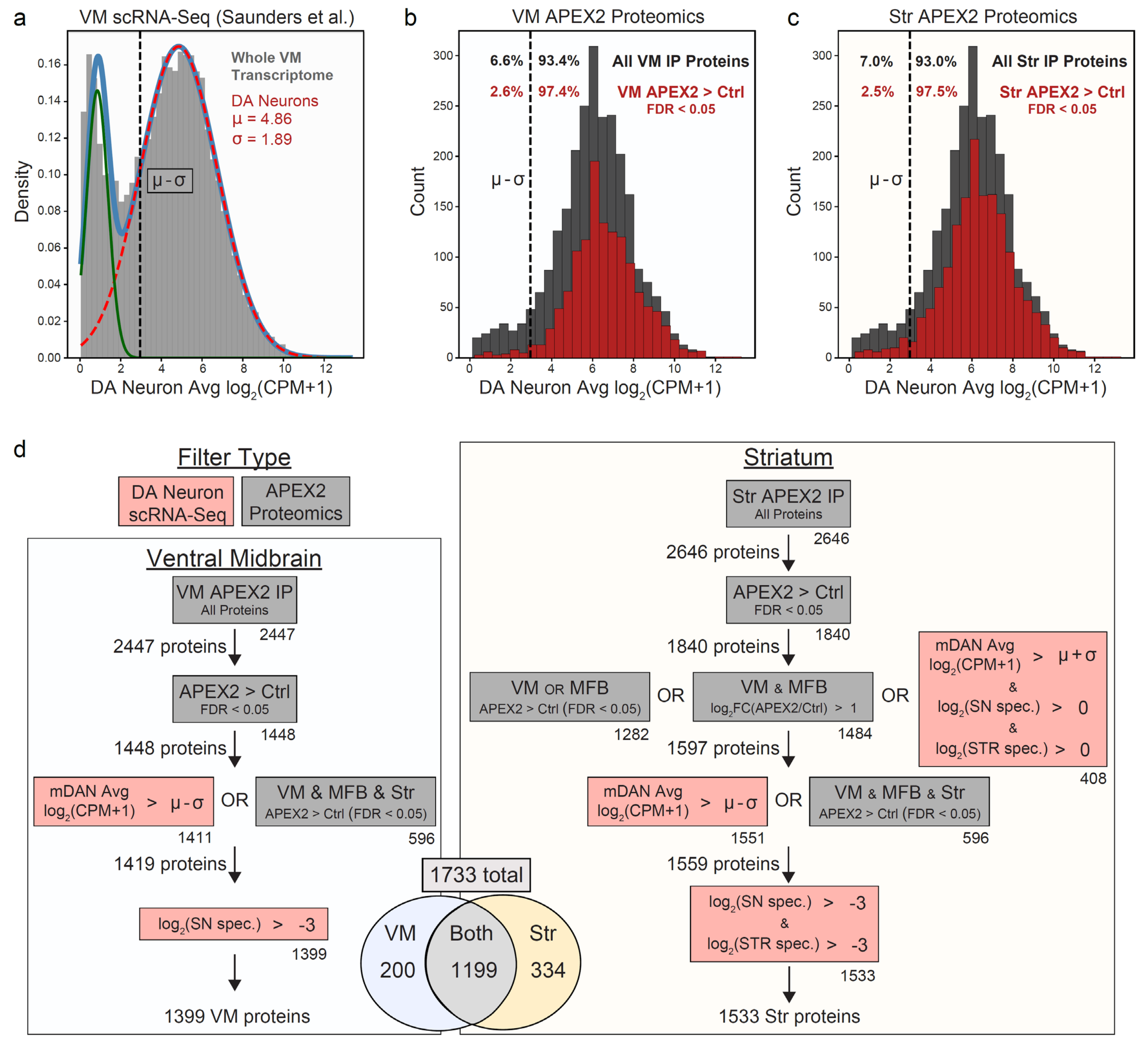


**Figure 3 – figure supplement 1: Filtering of APEX2 proteomics data using single-cell RNA sequencing (sc-RNAseq) data and cross-regional comparisons**

(**a**) Histogram of average mRNA expression in DA neurons identified in our reanalysis of the DropViz scRNA-Seq data (Saunders et al., 2018), see also **Figure 5 – figure supplement 1**. Data were fit with a gaussian mixture model (light blue trace), with the two gaussian distributions representing genes expressed (dashed red trace) or not expressed (green trace) in DA neurons. The mean (μ) and standard deviation (σ) for the DA neuronal component are indicated in red. A conservative threshold of μ - σ (dashed black line) was set as the lower bound to consider a gene expressed in DA neurons.

(**b**) Histogram of average DA neuron mRNA expression for all proteins detected in VM APEX2$^{+}$ IP samples (dark grey) and proteins enriched in VM APEX2$^{+}$ vs. APEX2$^{-}$ IP samples (FDR < 0.05, red). Only 6.6% of all VM IP proteins were below the lower bound of DA neuronal expression, which was decreased to 2.6% for proteins enriched in APEX2$^{+}$ vs. APEX2$^{-}$ IPs.

(**c**) Histogram of average DA neuron mRNA expression for all proteins detected in Str APEX2$^{+}$ IP samples (dark grey) and proteins enriched in Str APEX2$^{+}$ vs. APEX2$^{-}$ IP samples (FDR < 0.05, red). Only 7.0% of all Str

IP proteins were below the lower bound of DA neuronal expression, which was decreased to 2.5% for proteins enriched in APEX2$^+$ vs. APEX2$^-$ IPs.

(**d**) Filtering strategy for VM (*left*) and striatum (*right*) APEX2 proteomics data. Filters using scRNA-Seq data are shown in red, while filters using the proteomics data are shown in grey. Numbers below the bottom right corner of each box indicate the number of proteins passing that individual filter, while the total number of proteins passing each filter set is indicated to the left of the arrows. See **Methods** for detailed description of the filters. In total, 1399 and 1533 proteins were retained from the VM and striatum, respectively, with 1733 total. For most downstream analyses, the union of VM and striatum filtered proteins (1733) are referred to as the filtered APEX2 proteomics data. See **Figure 3 – source data 2** for complete results and summary of proteins before and after filtering.

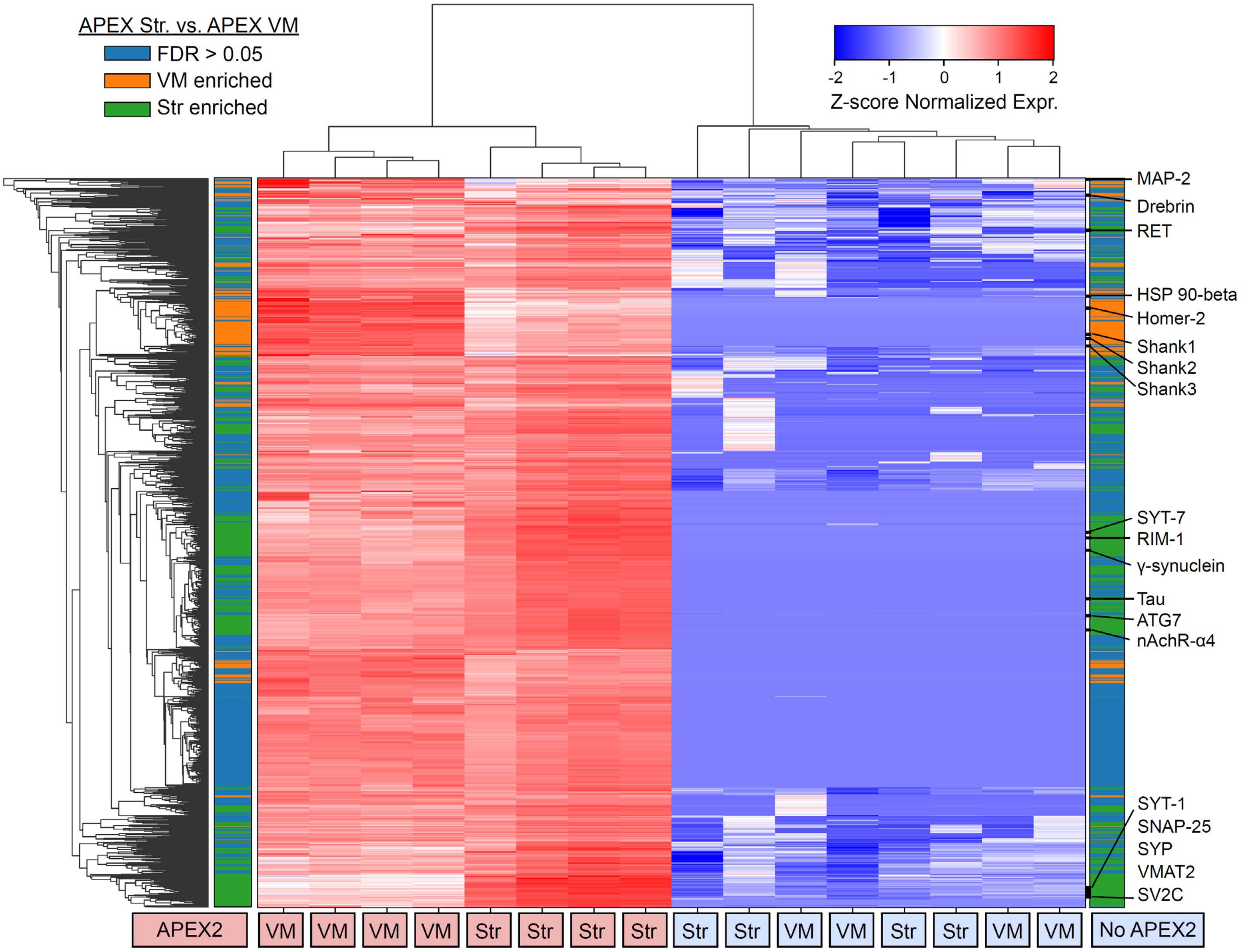


**Figure 3 – figure supplement 2**: **Clustered heatmap of protein abundances for union of filtered VM and Str APEX2 proteins**

Clustered heatmap of Z-scores for abundances of proteins present in the filtered proteomics data from VM or striatum (1733 proteins). Each column represents a biological replicate (n=4) of APEX2$^+$ or APEX2$^-$ streptavidin IP samples from the VM or striatum. The color bars on the left and right are identical; both indicate whether a given protein was enriched in the VM (orange) or striatum (green) in differential expression analysis between APEX2$^+$ IP samples (FDR < 0.05 after Benjamini-Hochberg corrected p-values from Welch's t-test). Select proteins from within given VM- or Str-enriched protein clusters are displayed on the right.

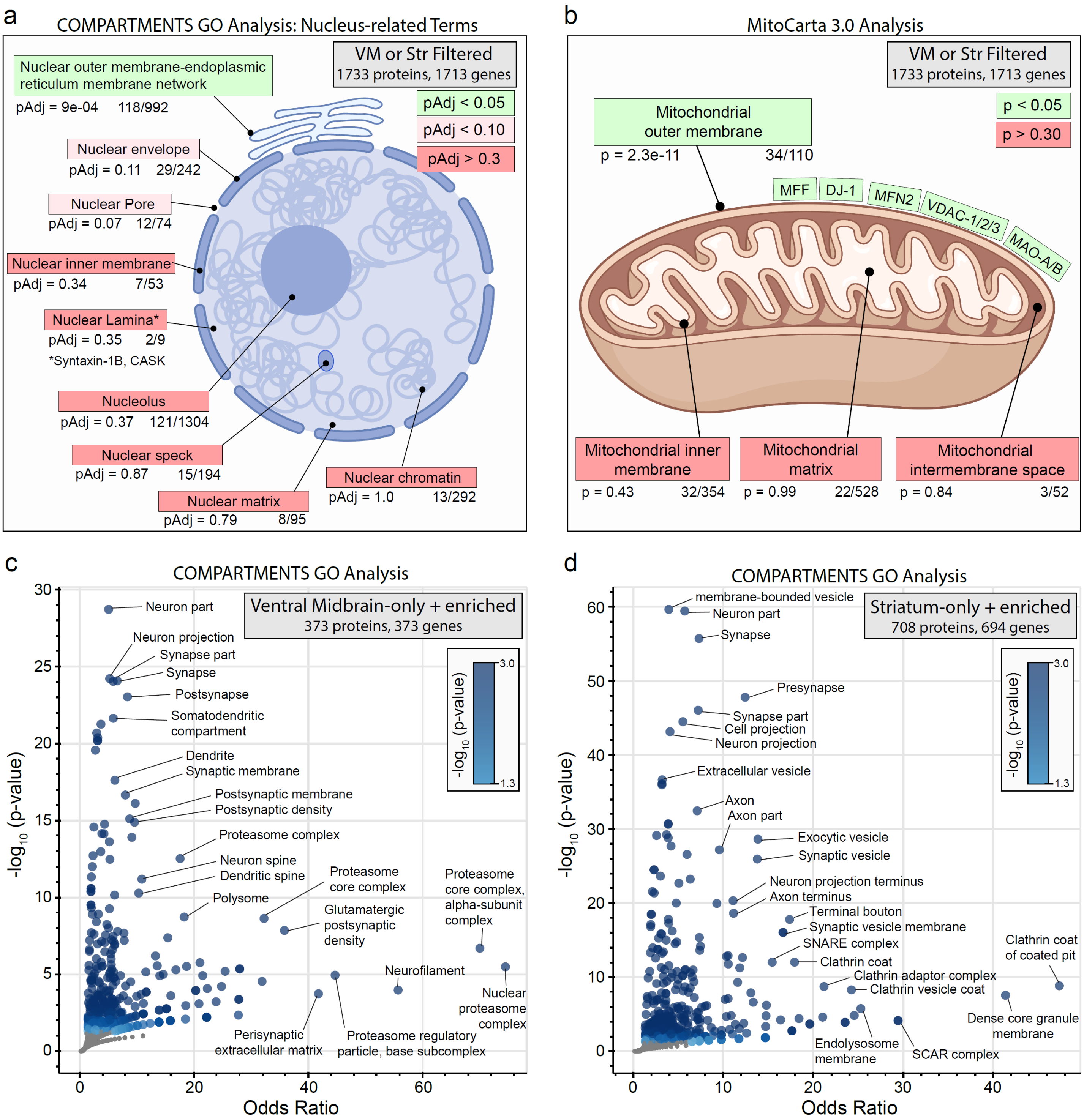


**Figure 3 – figure supplement 3**: **Subcellular compartment gene ontology analysis of filtered VM and Str APEX2 proteins**

(**a**) Targeted gene ontology analysis of proteins in filtered VM or striatum APEX2$^{+}$ data (1733 proteins encoded by 1713 genes). Proteins were analyzed for overlap with GO terms related to nucleus in the COMPARTMENTS resource (Binder et al., 2014). The number of proteins present in the filtered APEX2 data out of all proteins in each ontology is shown below each term, with FDR-corrected p-values derived from the hypergeometric test.

(**b**) Same as (**a**) but using mitochondrial compartmental localizations present in the MitoCarta 3.0 database (Rath et al., 2021). Select proteins present in the mitochondrial outer membrane list are displayed.

(**c**) Enrichr-based gene ontology analysis of proteins passing filter only in VM APEX2$^+$ samples or enriched in APEX2$^+$ VM vs. striatum differential expression (in total, 373 genes encoding 373 proteins). The 373 genes were analyzed using the subcellular compartments ontology terms provided by the COMPARTMENTS resource (Binder et al., 2014). All GO terms with $p < 0.05$ are colored in blue, with select terms indicated for those displaying the lowest p-values and highest odds ratio. See **Figure 3 – source data 3** for complete results.

(**d**) Same as (**c**) but for proteins passing filter only in striatum APEX2$^+$ samples or enriched in APEX2$^+$ striatum vs. VM differential expression (in total, 694 genes encoding 708 proteins).

a

Str-Slice / Syn Filtered
1348 proteins, 1329 genes → 393 SynGO annotations

| Rank | Ontology Term (Cellular Component) | Gene count | pAdj |
|---|---|---|---|
| 1 | synapse | 353 | 3e-110 |
| 2 | presynapse | 197 | 7e-66 |
| 3 | postsynapse | 179 | 1e-46 |
| 4 | synaptic vesicle | 63 | 2e-27 |
| 5 | synaptic vesicle membrane | 54 | 1e-23 |
| 6 | postsynaptic specialization | 82 | 3e-19 |
| 7 | postsynaptic density | 71 | 4e-48 |
| 8 | presynaptic active zone | 33 | 2e-11 |
| 9 | presynaptic membrane | 38 | 6e-10 |
| 10 | presynaptic cytosol | 19 | 2e-09 |

| Rank | Ontology Term (Biological Process) | Gene count | pAdj |
|---|---|---|---|
| 1 | process in the synapse | 283 | 4e-85 |
| 2 | process in the presynapse | 124 | 3e-48 |
| 3 | synaptic vesicle cycle | 102 | 2e-44 |
| 4 | synapse organization | 96 | 5e-27 |
| 5 | process in the postsynapse | 68 | 9e-19 |
| 6 | synaptic vesicle exocytosis | 40 | 1e-17 |
| 7 | synaptic vesicle endocytosis | 32 | 2e-15 |
| 8 | regulation of postsynaptic membrane neurotransmitter receptor levels | 45 | 6e-15 |
| 9 | trans-synaptic signaling | 54 | 2e-14 |
| 10 | chemical synaptic transmission | 49 | 9e-14 |

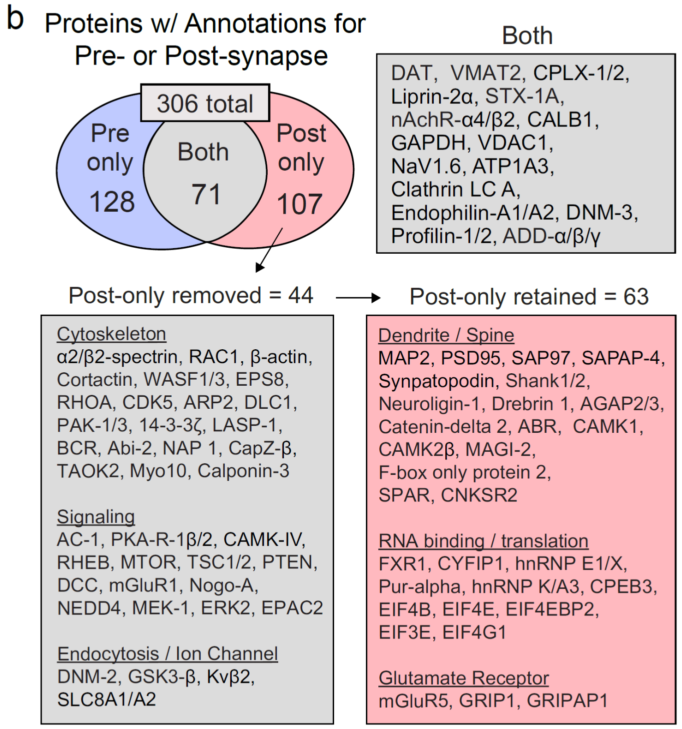


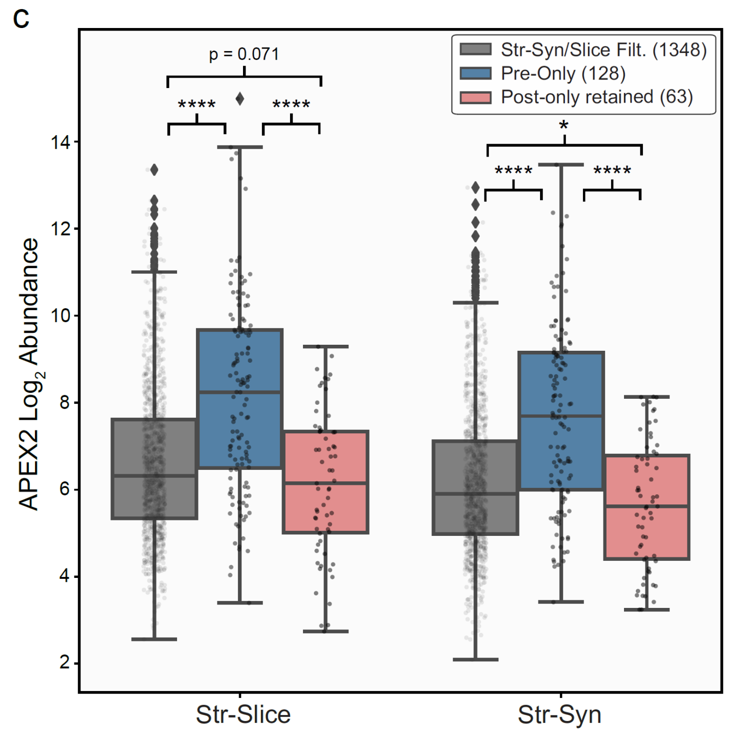


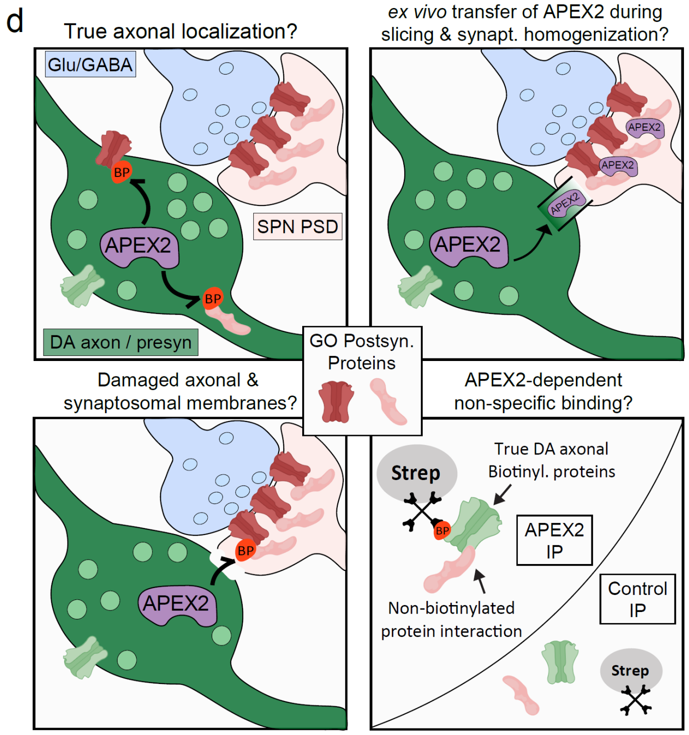


**Figure 4 – figure supplement 1**: **Synaptic gene ontology analysis proteins of striatal slice and synaptosomal APEX2 data**

(**a**) SynGO (Koopmans et al., 2019) analysis of striatum slice and synaptosome filtered proteins (1348 proteins encoded by 1329 genes, with 393 SynGO annotations). The top 10 Ontology Terms for Cellular Component and Biological Process are shown. See **Figure 4 – source data 2** for complete results.

(**b**) Downstream filtering of 179 genes from Cellular Component term “postsynapse”. 71 proteins with dual annotation to presynapse and postsynapse were removed from the “postsynapse” set (proteins such as DAT, VMAT2, etc., shown in “Both”). The remaining 107 proteins were designated as having “Post-only” annotations, and were further filtered to remove generic cytoskeletal, signaling, and synaptic proteins with functions in both pre- and post-synapse. All 44 proteins removed are shown under “Post-only removed”. The remaining 63 proteins were designated as "Post-only retained" for downstream analysis. See **Figure 4 – source data 3** for complete list of protein abbreviations in (**b**).

(**c**) APEX2 samples average $\log_2$(normalized intensity + 1) for indicated protein sets derived from SynGO analysis (panels **a-b**). * indicates p < 0.05 and **** indicates p < 0.0001 from Mann-Whitney U test. Mann-Whitney test statistics are as follows, *left to right*: Str-Slice Pre vs. All ($U$ = 40948, p = 1.4e-14), Str-Slice Post vs. All ($U$ = 30448, p = 0.071), Str-Slice Pre vs. Post ($U$ = 1986, p = 6.2e-09), Str-Syn Post vs. All ($U$ = 29166, p = 0.024), Str-Syn Pre vs. All ($U$ = 39841, p = 1.3e-15), Str-Syn Pre vs. Post ($U$ = 1793, p = 2.3e-10).

(**d**) Schematic depicting possible mechanisms underlying striatal APEX2 enrichment of proteins associated with classical postsynaptic gene ontologies ("Post-only retained" shown in **b-c**).

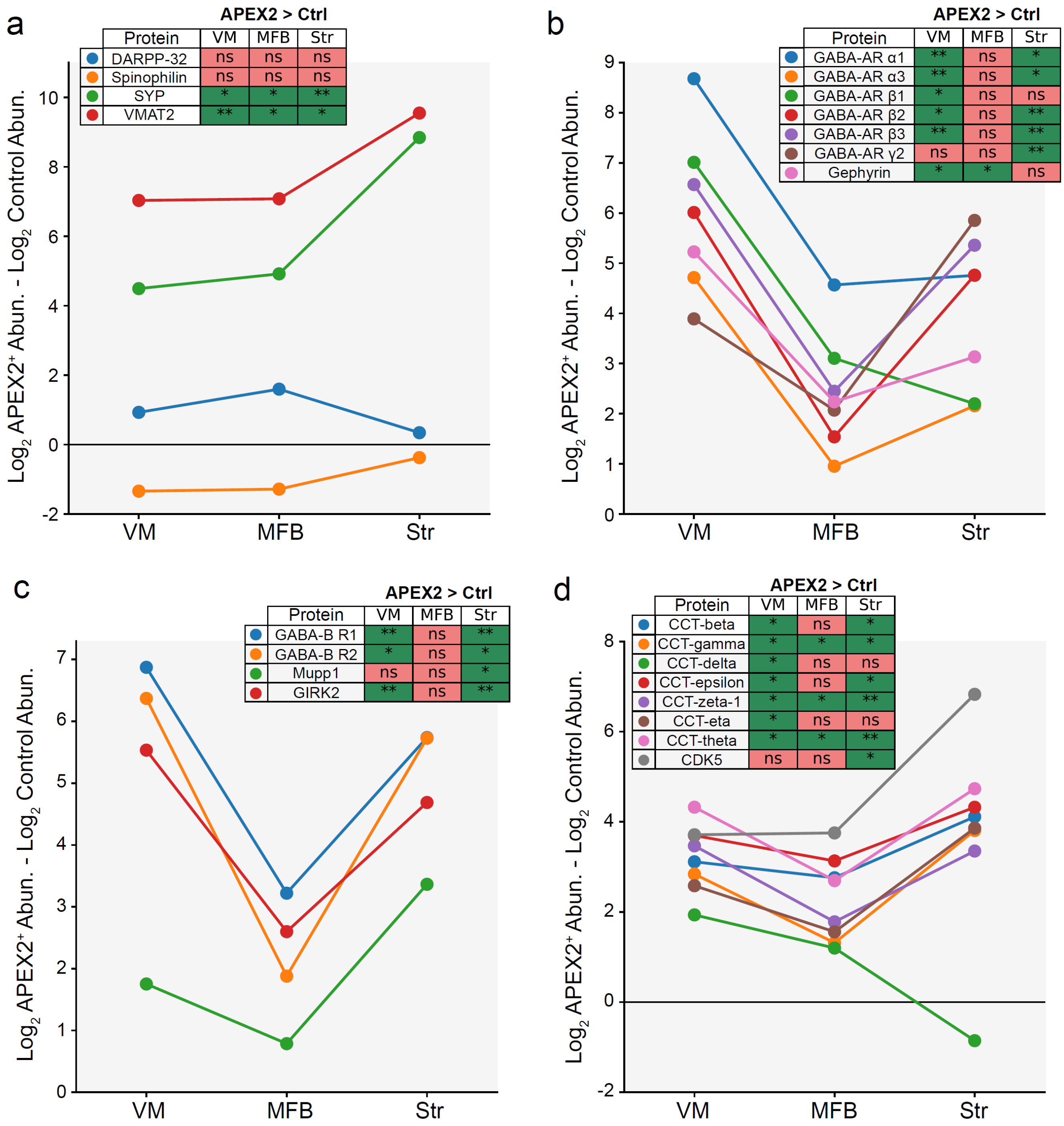


**Figure 4 – figure supplement 2**: **APEX2 enrichment of specific protein groups across DA neuronal regions**

For all panels, the difference in average $\log_2$(normalized intensity + 1) between APEX2$^+$ and APEX2$^-$ (control) samples is plotted for the indicated proteins. The legend indicates the result of the Welch's unequal variance t-test with Benjamini-Hochberg procedure to control the FDR (n = 4 biological replicates each for APEX2$^+$ and APEX2$^-$ samples in each region). * indicates FDR < 0.05, ** indicates FDR < 0.001.

**(a)** Dendritic spine proteins DARPP-32 and Spinophilin are not enriched in APEX2 striatal samples, while pre-synaptic proteins Synaptophysin and VMAT2 are massively enriched.

**(b)** GABA-A receptor subunits and scaffolding protein Gephyrin are strongly enriched by APEX2 in the ventral midbrain, but are also captured in the medial forebrain bundle and striatum.

**(c)** GABA-B receptor subunits, PDZ-domain containing scaffolding protein Mupp1, and effector ion channel GIRK2 are captured by APEX2 in both VM and striatum.

**(d)** CDK5 and most all members of the eukaryotic group II chaperonin TRiC (tailless complex polypeptide 1 ring complex) are captured by APEX2 in the VM and striatum.

*Protein Abbreviations*: (DARPP-32) dopamine- and cyclic-AMP-regulated phosphoprotein of molecular weight 32 kDa, (SYP) synaptophysin, (VMAT2) vesicular monoamine transporter 2, (GABA-AR) GABA-A receptor subunit, (GABA-BR) GABA-B receptor subunit, (GIRK2) G protein-activated inward rectifier potassium channel 2, (CCT) chaperonin-containing tailless complex polypeptide 1 subunit, (CDK5) cyclin dependent kinase 5

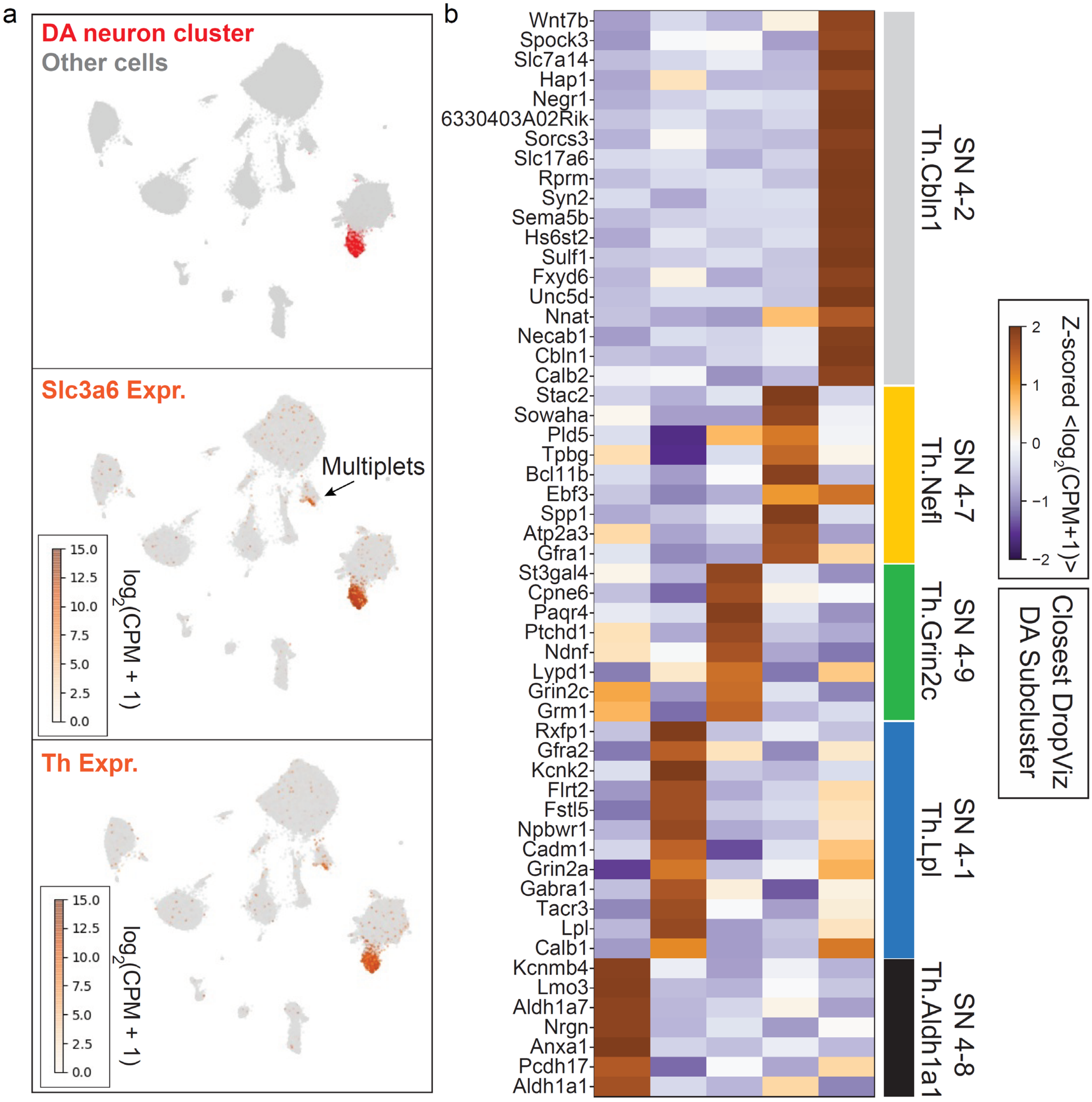


**Figure 5 – figure supplement 1**: **Identification of DA neuron cluster and sub-clusters for downstream analysis**

(**a**) UMAP embedding of scRNA-Seq data from (Saunders et al., 2018). A single cluster with statistically significant co-enrichment of dopamine neuron markers such as *Th* and *Slc6a3*, based on the binomial test for expression specificity, was identified. After sub-clustering the putative dopamine neurons, we identified a small sub-cluster with statistical enrichment of astrocyte markers such as *Agt*, *Gja1*, *Glul*, and *Slc1a3*. for pan-DA neuronal expression analysis. This sub-cluster was discarded due to likely astrocyte contamination. After removal of low-quality cells, we retained the remaining subclusters as high-confidence DA neuron profiles (see **Methods**). DA neuron profiles used in this study are found in **Figure 5 – source data 3**.

(**b**) Sub-clustering of high-confidence DA neurons identified five transcriptionally distinct DA neuron subsets. Markers determined using the binomial test are shown in the heatmap, and were used to identify the closest corresponding cluster present in the DropViz data (Saunders et al., 2018).

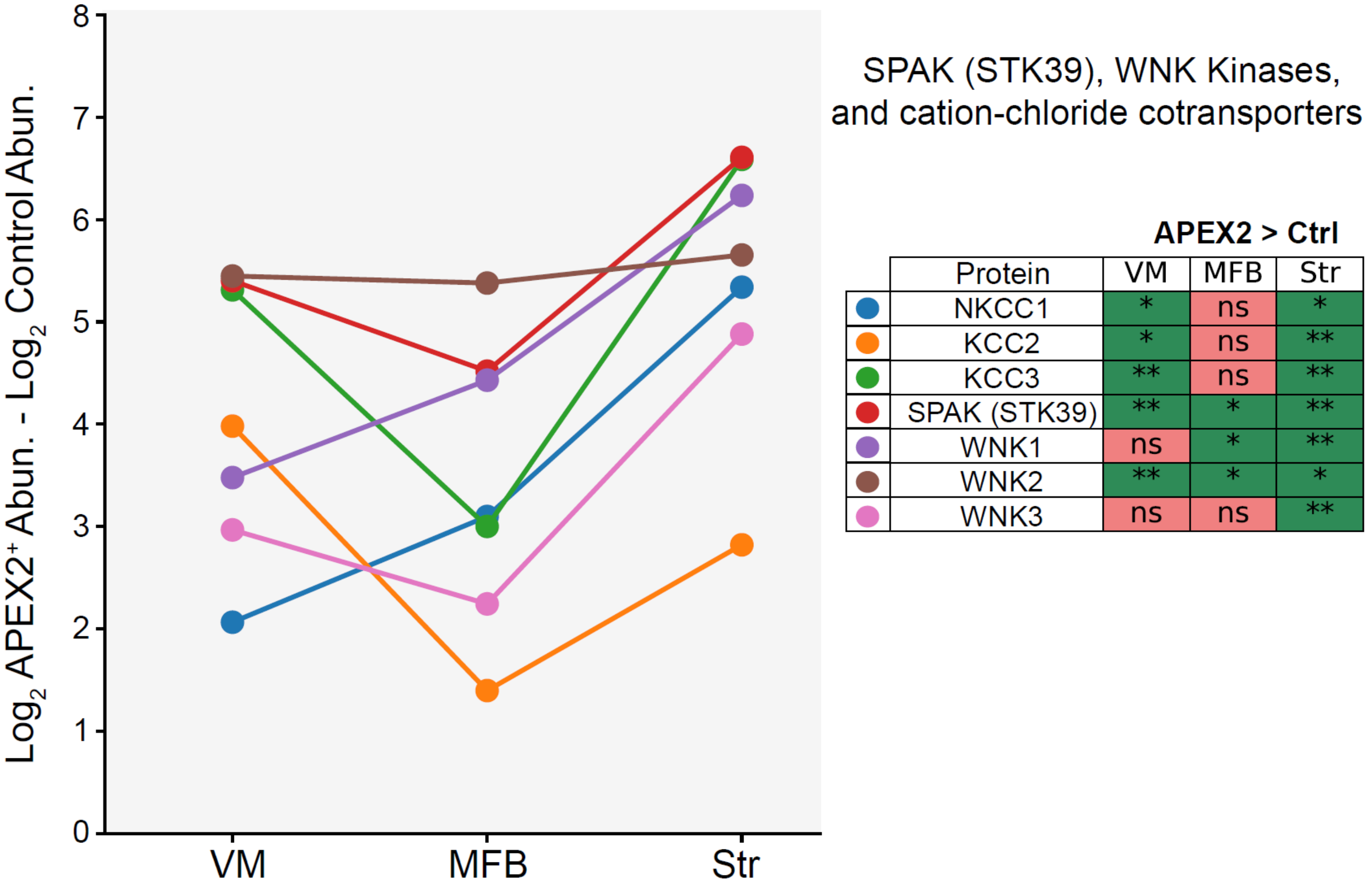


| Protein | VM | MFB | Str |
|---|---|---|---|
| NKCC1 | * | ns | * |
| KCC2 | * | ns | ** |
| KCC3 | ** | ns | ** |
| SPAK (STK39) | ** | * | ** |
| WNK1 | ns | * | ** |
| WNK2 | ** | * | * |
| WNK3 | ns | ns | ** |

**Figure 5 – figure supplement 2**: **SPAK and associated proteins are present in DA axons**

The difference in average log$_2$(normalized intensity + 1) between APEX2$^+$ and APEX2$^-$ (control) samples is plotted for the indicated proteins. The legend indicates the result of the Welch's unequal variance t-test with Benjamini-Hochberg procedure to control the FDR (n = 4 biological replicates each for APEX2$^+$ and APEX2$^-$ samples in each region). * indicates FDR < 0.05, ** indicates FDR < 0.001.

*Protein Abbreviations*: (NKCC1) Solute carrier family 12 member 2 / basolateral Na-K-Cl symporter, (KCC2) Solute carrier family 12 member 5 / K-Cl cotransporter 2 / mKCC2, (KCC3) Solute carrier family 12 member 6 / K-Cl cotransporter 3, (SPAK / STK39) STE20/SPS1-related proline-alanine-rich protein kinase / Ste-20-related kinase, (WNK) Serine/threonine-protein kinase WNK / Protein kinase with no lysine
